# Tandem MutSβ binding to long extruded DNA trinucleotide repeats underpins pathogenic expansions

**DOI:** 10.1101/2023.12.12.571350

**Authors:** Jun Li, Huaibin Wang, Wei Yang

**Affiliations:** Laboratory of Molecular Biology, NIDDK, National Institutes of Health, Bethesda, MD 20892; Laboratory of Cell and Molecular Biology, NIDDK, National Institutes of Health, Bethesda, MD 20892

**Keywords:** repeat expansion, neurological disorders, mismatch repair, insertion deletion loop (IDL)

## Abstract

Expansion of trinucleotide repeats causes Huntington’s disease, Fragile X syndrome and over twenty other monogenic disorders^1^. How mismatch repair protein MutSβ and large repeats of CNG (N=A, T, C or G) cooperate to drive the expansion is poorly understood. Contrary to expectations, we find that MutSβ prefers to bind the stem of an extruded (CNG) hairpin rather than the hairpin end or hairpin-duplex junction. Structural analyses reveal that in the presence of MutSβ, CNG repeats with N:N mismatches adopt a B form-like pseudo-duplex, with one or two CNG repeats slipped out forming uneven bubbles that partly mimic insertion-deletion loops of mismatched DNA^2^. When the extruded hairpin exceeds 40-45 repeats, it can be bound by three or more MutSβ molecules, which are resistant to ATP-dependent dissociation. We envision that such MutSβ-CNG complexes recruit MutLγ endonuclease to nick DNA and initiate the repeat expansion process^3,4^. To develop drugs against the expansion diseases, we have identified lead compounds that prevent MutSβ binding to CNG repeats but not to mismatched DNA.

## Introduction

DNA sequence contractions and expansions often occur at microsatellites of 3-6 nucleotide (nt) repeats^5,6^. Expansions of gene-specific CNG (N=A, C, G or T) trinucleotide repeats (TNRs) alone cause 26 monogenic and often neurological disorders including Huntington’s disease (HD), fragile X syndrome (FXS), myotonic dystrophy type 1 (DM1), and many spinocerebellar ataxias (SCAs)^1^. In parallel, genome-wide instability at microsatellites of 1-2 nt repeats due to replication slippage and mismatch repair deficiency is a hallmark of colorectal and Lynch syndrome cancers^7,8^. These microsatellites, which account for 3% of the human genome^9,10^ are important for genome regulation and diversity but are polymorphic and unstable. Paradoxically among mismatch repair (MMR) proteins, which normally maintain microsatellite stability by removing mis-incorporated nucleotides and insertion-deletion loops (IDLs) formed during DNA replication^11,12^, MutSβ (MSH2-MSH3 heterodimer) and MutLγ (MLH1-MLH3 dimer) are drivers of CNG and other TNR instability^1,4,6,13–15^. TNR expansions are repeat length dependent. Below the threshold of ∼40 repeats TNRs are stable, but beyond that, the probability of expansion increases rapidly in correlation with the size of repeats^16,17^. In addition, repeat lengths are often correlated with the age of onset, progression and the severity of repeat-expansion diseases^18^. The requirement for both long repeats and MMR proteins suggests a conserved mechanism for TNR expansions.

CNG repeats slipped out from DNA double helix (slip-outs or S-DNA) have been observed to form hairpins, in which every two successive C/G basepairs are followed by an N:N mismatch^19–22^ (Fig. 1a). While no one has examined if MutSβ binds to such a mismatched hairpin without surrounding basepaired duplex DNA, MutSβ was shown to bind a (CNG)_13_ (13 repeats) sequence embedded in a DNA duplex with K_d_ of ∼5 nM^23,24^. Because MutSβ binds to IDLs of 2-14 nt embedded in homoduplex and bends the DNA duplex 80-110° at the IDL^2^, interacting with the surrounding homoduplex is essential for mismatch recognition. The slipped-out CNG repeats was hypothesized to partially mimic IDL but trap MutSβ at the hairpin ends or duplex-hairpin junctions and prevent MMR by altering the MutS ATPase activity^23,25^. However, the efficient removal of (CNG)_n_ slip-outs (n ≤ 25) by MMR and other repair pathways^26,27^ contest this hypothesis.

**Fig. 1.**
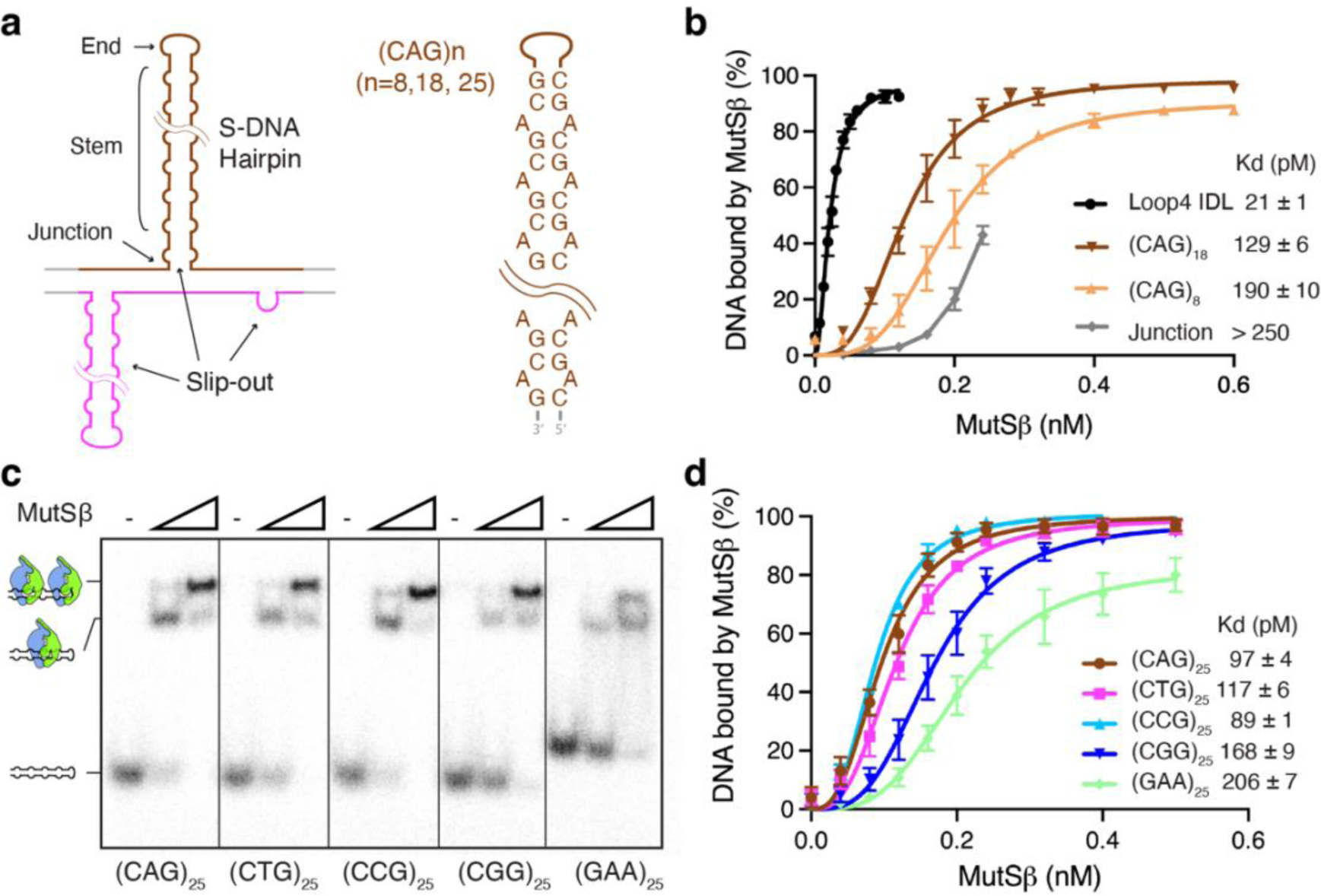
MutSβ binding to CNG repeats. **a,** Diagrams of S-DNA formed at a CNG locus (left) and (CAG)n hairpins (n=8, 18 or 25) (right) without surrounding homoduplex. **b,** K_d_ plot of MutSβ binding to the hairpin end ((CAG)_8_), hairpin stem ((CAG)_18_), hairpin junction (TriArm_9-9), and insertion-<u>d</u>eletion loop (Loop4 IDL) based on triplicate EMSA measurements. **c-d,** EMSA analysis (**c**) and K_d_ plot (**d**) of MutSβ binding to four variations of (CNG)_25_ and (GAA)_25_.

In the absence of DNA replication, the CAG repeats in the coding region of Huntingtin gene of HD patients are expanded in non-dividing neurons^28^, and age-dependent repeat expansions have also been observed in non-mitotic cells in mouse models^29^. Repeat expansions without replication require DNA nicking or a double-strand break to precede repair-coupled synthesis and expansion. Repeat expansion formed in multiple steps, involving initial DNA cleavage and following repair synthesis, has been suggested^30,31^. Previously, in addition to nucleases cleaving at oxidized bases (OGG1) or unusual DNA structures (FAN1) being implicated in repeat instability^3,32^, recognition of stably engineered S-CNG by MutSβ has been shown to activate MutLα (MLH1-PMS2 dimer) and MutLγ to nick DNA *in vitro*^4,33^. But of the two, only MutLγ is required *in vivo* for CAG repeat expansion in Huntington’s Disease^34^. After initial DNA nicking, MutSβ may stabilize misaligned template and primer strands of CNG repeats and cause deletions or insertions of one-repeat unit during repair DNA synthesis^35^.

A long-standing question is how S-CNG hairpins, which are less stable than the alternative homoduplex and readily dissociated from MutSβ due to the MutSβ ATPase activity^27,36–38^, stay bound to MutSβ and lead to DNA nicking and repeat expansions. We report here binding properties and cryoEM structures of (CNG)_n_ hairpins (n= 8 to 60, both below and beyond the threshold of expansion) complexed with human MutSβ, which shed light on the initial steps in the accelerated CNG expansion and pathogenesis with large repeats.

## Results

### MutSβ prefers to bind the stems of (CNG)n hairpins

To find out if and where MutSβ may bind CNG repeat hairpins without surrounding DNA, we prepared 8 and 18 CAG repeats ((CAG)_8_ and (CAG)_18_), which potentially form 11 and 26 bp heteroduplex hairpins with 2-nt turns (Fig. 1a). As every MutS homolog, human MutSβ included, covers 20-25 bp of DNA duplex with mismatched nucleotides in the middle, (CAG)_8_ presents half a binding site and only the hairpin end (turn) may be bound by MutSβ, while (CAG)_18_ may be bound by MutSβ either at hairpin end or the middle (stem). Using picomolar (pM) DNA concentrations without competitor DNAs in EMSA (electrophoretic mobility shift assay) analysis^2^, we measured the K_d_ values in a greater dynamic range and 50-100 fold lower than reported in other studies^23,25,33^. Human MutSβ bound 10 pM of these ^32^P-labeled CAG repeats and formed a single protein-DNA complex in each case with dissociation constant (K_d_) of 190 pM and 129 pM for the (CAG)_8_ and (CAG)_18_ hairpins, respectively (Fig. 1b, Extended Data Fig. 1a). Although binding to the CAG hairpins is weaker than the Loop4-IDL (4-nt insertion) DNA (K_d_=21 pM) (Fig. 1b), the EMSA analyses indicated that MutSβ can bind the hairpin stem as well as the end and binding to the stem ((CAG)_18_) is tighter than the hairpin end ((CAG)_8_).

We then prepared (CAG)_25_, which can form a 36 bp heteroduplex hairpin assuming 3 nt at the hairpin end and is long enough to accommodate two MutSβ molecules, and found two distinctly MutSβ-shifted DNA bands (Fig. 1c). Based on the binding analyses of (CAG)_8_ and (CAG)_18_, the first band is likely a result of MutSβ binding to the stem of the CAG hairpin, and the second a result of two MutSβ molecules binding simultaneously to the stem and the hairpin end. EMSA of (CTG)_25_, (CCG)_25_, and (CGG)_25_ with MutSβ is consistent with the protein binding to both the stem and end of (CNG)_25_ hairpins, with two-site combined K_d_ ranging from 89 (CCG) to 168 pM (CGG) (Fig. 1c-d, Extended Data Fig. 1b). Even (GAA)_25_, whose expansion causes Friedreich’s Ataxia^39^, is bound by MutSβ with K_d_ of 206 pM in the absence of any Watson-Crick basepair. In contrast, binding to the junction, where a (CAG)n hairpin extrudes from DNA duplex, is weaker (Fig. 1a-b, Extended Data Fig. 1c-d). The weak and flexible junction binding by MutSβ was confirmed by cryoEM (Extended Data Fig. 1e).

To verify (CAG)n pseudo-duplex binding by MutSβ, we prepared an 80 bp duplex with a (CAG)_30_ inserted in the middle of one strand (d40h30) and visualized its association with MutSβ by cryoEM. Preliminary particle analysis revealed that 78% of MutSβ bound to the hairpin stem, 22% to the hairpin end, and none to the junction at the 2:1 molar ratio of protein and DNA (Extended Data Fig. 2a, Methods). Combined EMSA and cryoEM analyses show that the stem of an extruded (CAG)n hairpin is the primary site for MutSβ binding, the hairpin end is the second, and the S-DNA junction the last.

### MutSβ binds (CAG)_25_ and (CTG)_25_ hairpin stems as IDLs

To find out how MutSβ binds CNG heteroduplexes, we determined the structures of MutSβ complexed with (CAG)_30_ (d40h30) as well as (CAG)_25_ by cryoEM. The two samples were indistinguishable, and the data were combined. As binding to the stems and ends of CAG hairpins co-existed, these structures were separately classified and refined to 2.8 to 3.4 Å resolution, respectively (Extended Data Fig. 2 and Table 1). The protein-stem complexes contain one or two CAG <u>u</u>nit <u>s</u>lip-out (1-us or 2-us, or insertion of 3 or 6 nt) in the middle of the heteroduplex, forming uneven DNA bubbles with 5 or 8 nt on one side opposite 2 nt on the other (see detail below), and the 1-us is dominant (>90%) (Extended Data Fig. 2a). The stem-binding MutSβ-(CAG)_25_ structure is largely indistinguishable from the crystal structures of MutSβ-IDL complexes^2^ (Fig. 2a-c). The complex of MutSβ and hairpin-end is also superimposable except for missing the half of DNA duplex after the hairpin end (Fig. 2b, Extended Data Fig. 2j). Of the potential heteroduplex, 21 bp can be modeled as a double helix, 8 and 13 bp on each side of the ∼90° DNA bend at the slip-out (Extended Data Fig. 3a). The DNA duplex upstream (5′) of the insertion extensively interacts with the mismatch recognition domain I of MSH3 (M3_I) and DNA binding domain IV of MSH2 (M2_IV)^2^ and is thus well ordered and termed the “stable arm” (Fig. 2c). The duplex downstream (3′) of the insertion is termed the “mobile arm” because it is flexible and less well defined due to limited interactions with MutSβ^2^. The well-resolved cryoEM map of the stable arm reveals that MutSβ binds the phosphosugar backbones with the A:A mismatches forming B-like DNA (Fig. 2d and 3a). Two mismatched As adopt the *anti* and *syn* conformations as in Hoogsteen pairs except for only one hydrogen bond between them, and the C1′-C1′ distance is 10.5 Å, close to the 10.6 Å in B-DNA (Fig. 2e). This finding is related to but different from the previous foot-printing study of the half hairpin ((CAG)_13_) embedded in homoduplex^23^. Normally, a single mismatched base pair disrupts base stacking and caused the DNA duplex to flex, so it is recognized by MutS protein for mismatch repair^36,40^. It is stunning that MutSβ, which binds IDLs that are flanked by homoduplexes, can induce and stabilize the heteroduplex of CAG repeats into B-like DNA with every third base pair being an A:A mismatch!

**Fig. 2.**
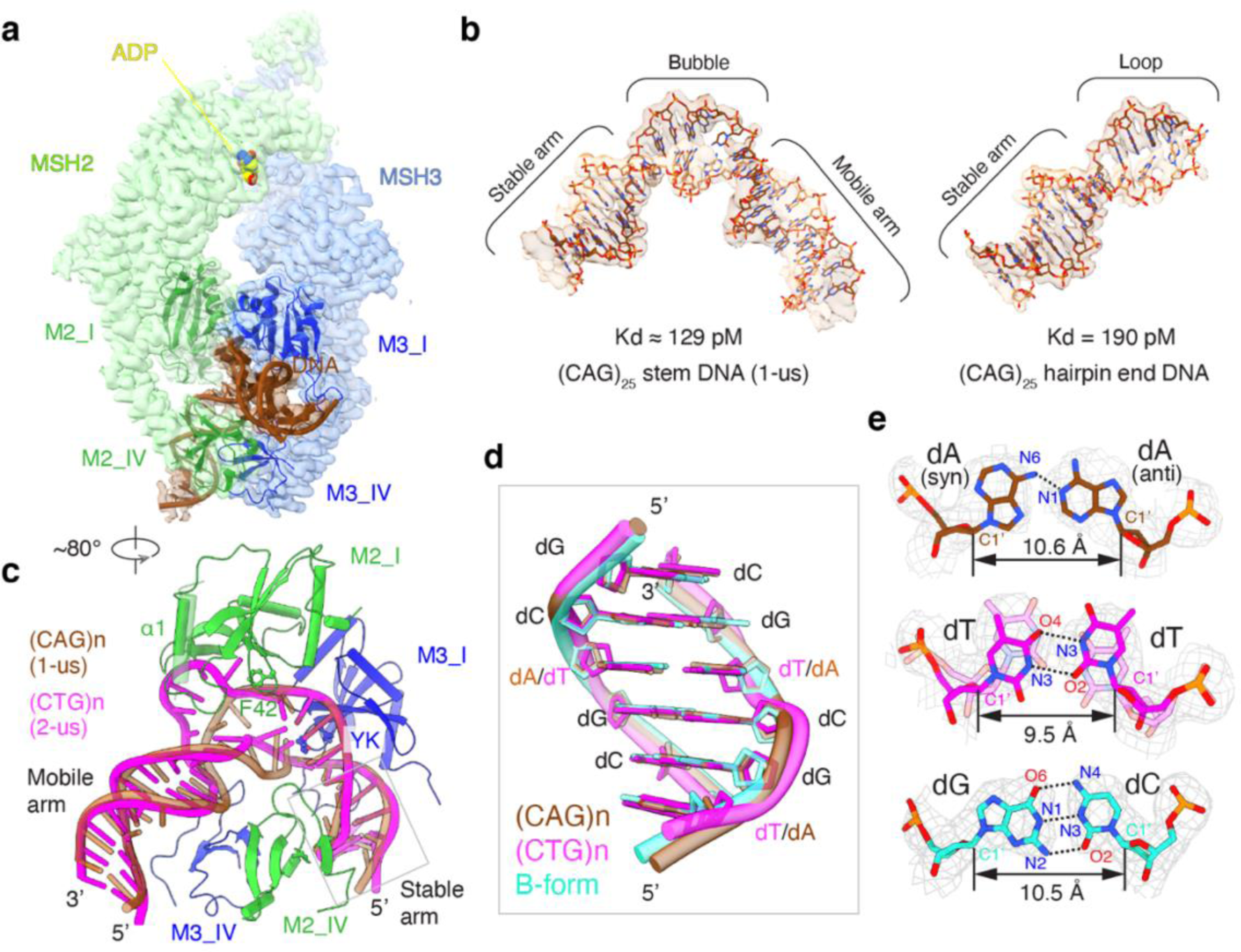
Structures of (CAG)_25_ and (CTG)_25_ hairpin in complex with MutSβ. **a,** CryoEM map of MutSβ-(CAG)_25_ hairpin stem complex. Cartoons of DNA and domains I and IV of MSH2 and MSH3 are superimposed. The ADP bound to MSH2 is also shown. **b,** Structures of the (CAG)_25_ hairpin stem and end. **c,** The kinked 1-us (CAG)n (brown) and 2-us (CTG)n (magenta) are stabilized by MSH2 (green) and MSH3 (blue). The longer α1 helix of M2_I in the (CAG)n structure is shown in pale green. The YK motif (MSH3) and F42 (MSH2) are shown as ball-and-sticks. **d,** A zoom-in view of DNA in the stable arm (boxed in panel **c**). CAG (brown) and CTG (magenta) pseudo-duplexes are superimposed with a B-form homoduplex (cyan). **e,** The mismatched A and A are in *syn* and *anti* configuration. The T:T mismatch adopts two isoforms. Dotted lines depict Hydrogen bonds. The C1′-C1′ distance is indicated.

**Fig. 3.**
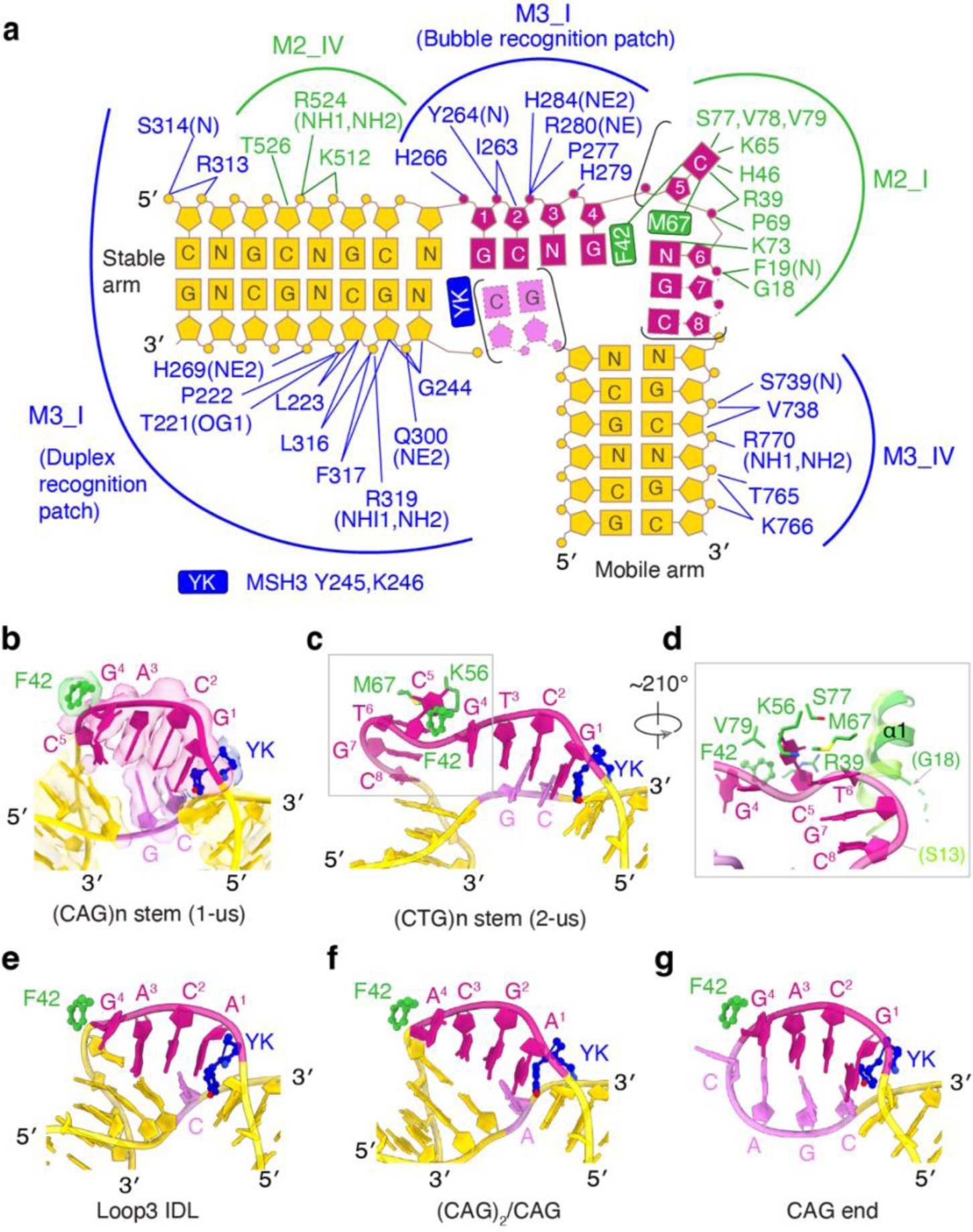
Recognition of uneven DNA bubbles. **a,** Diagram of MutSβ and (CNG)n (N=A or T) pseudo-duplex interactions. **b-c,** The 1-us (CAG)n (**b**) and 2-us (CTG)n bubble (**c**) are framed by the YK motif of MSH3 (blue) and Y42 of MSH2 (green). The long and short side of uneven bubbles are colored dark and light pink, respectively, and the surrounding pseudo-duplex is colored gold. The cryoEM map (semi-transparent) is superimposed in panel **b**. K56 and M67 (MSH2) that stabilize the flipped-out base (C^5^) are shown as sticks (**c**). **d,** Detailed interactions between M2_I and the DNA bulge (C^5^ to C^8^) in the 2-us (CTG)n structure. Without uncoiling, the α1 helix of M2_I (semi-transparent cartoon) would clash with the DNA. **e-g,** The IDL structures of Loop3 (PDB: 3THX) (**e**), the uneven bubble of (CAG)_2_/CAG (**f**), and the hairpin loop of (CAG)n (**g**) in the same color scheme as (**b**) and (**c**).

Structures of MutSβ complexed with (CTG)_25_ were also determined (Extended Data Fig. 2 and Table 1). Unsurprisingly, MutSβ binds either the hairpin stem or end. Each T:T mismatch is formed by two hydrogen bonds in alternative pairing isomers (Fig. 2e). Even though the resulting C1′-C1′ distance is 9.5 Å, narrower than in B-DNA, the mismatches sandwiched between G:C basepairs are bound by MutSβ like homoduplex (Fig. 2d). Similar to (CAG)_25_, two different stem-binding structures with one or two CTG-units slipped-out co-existed. However, 2-us of (CTG)_25_ is twice as populated as 1-us, indicating a base composition-dependent unit slippage preference.

### Recognition of uneven bubbles formed by CNG repeats

To better characterize MutSβ binding to slip-out uneven DNA bubbles, we prepared DNAs with (CAG)_2_/CAG or (CAG)_2_/T_3_ inserted in the middle of a 40 bp duplex. EMSA analysis revealed that when flanked by homoduplex DNA, (CAG)_2_/CAG and (CAG)_2_/T_3_ uneven bubbles are bound by MutSβ with K_d_ of 56-66 pM (Extended Data Fig. 4a-b), ∼2-fold tighter than the (CAG)_25_ hairpin stem, which indicates the potential energy cost of molding A/A mismatches into B-like DNA.

Structures of these DNAs bound by MutSβ were well defined in cryoEM maps (Extended Data Fig. 4 and Table 1). The long side of the uneven bubble always contains a 4 nt insertion sandwiched between Y245-K246 of MSH3 (YK motif) and F42 of MSH2, like the extruded loops of IDLs^2^ (Fig. 3b-f). YK is intercalated between the paired (pseudo) duplex and unpaired bases at the 5′ end of the insertion, and F42 is π stacked with the fourth base of the insertion. Y245 forms a π-π stacking interaction with the single unpaired base opposite the 4-nt insertion. This 4:1 uneven bubble is observed in the (CAG)_2_/CAG complex (Fig. 3f). In the (CAG)_2_/T_3_, as well as 1-us CAG and CTG structures, the first basepair 3′ to the insertion is unpaired, which results in a 5:2 uneven bubble (Fig. 3b, Extended Data Fig. 3b). The 5′ and 3′ basepairs surrounding the 5:2 uneven bubble are usually hydrogen bonded even if they are buckled or sheared mismatches. The 4-nt insertion between YK and F42 can be of any sequence because of the lack of base-specific interactions of Mutβ.

A previously unknown mechanism of MutSβ-IDL interactions is revealed in the 2-us structures, where the uneven bubble consists of 8 and 2 nt on each side with CAG and CTG alike. The double-sized uneven bubbles kink the DNA 10° more than the 1-us DNAs (Extended Data Fig. 3a). Despite the very different bubble size, they retain the 4-nt insertion defined by the YK motif (MSH3) and F42 (MSH2) (Fig. 3c). Different from the 4:1 and 5:2 uneven bubbles, 2 nt (3′-CG-5′) on the short side are perfectly base paired with GC at the 5′ end of the insertion. Thus, the upstream DNA duplex is extended 2 bp into the bubble after the YK interaction, which results in unwinding but not bending. The lack of protein-DNA interactions on the short side of the uneven bubble allows the flexibility of these nucleotides to be base paired or not (Fig. 3).

The last four nucleotides (5^th^ to 8^th^) in the insertion loop are bulged out into a groove formed entirely within domain I of MSH2. A sharp 90° bend occurs between the 5^th^ and the 6^th^ nucleotides separated by M67 (Fig. 3c-d). The 5^th^ base is sandwiched between K65-M67 and F42 and V79 of MSH2, while the 6^th^ to 8^th^ unpaired bases are neatly stacked with the downstream heteroduplex. To accommodate this DNA bulge, the N-terminal first α-helix of MSH2 is uncoiled by one helical turn (Fig. 2c and 3d). Two following β hairpin turns contribute R39, P69, K73, S77, in addition to residues mentioned above, to interact with the DNA bulge (Fig. 3a). The bulge-binding site is likely utilized when MutSβ binds and repairs IDLs larger than 6 nt^41^. The unique role of MSH2 in the DNA bulge binding provides an explanation for why its domain I is required for IDLs binding by MutSβ but is dispensable in mispaired base recognition by MutSα^42^.

### MutSβ binding to hairpin ends and other TNR

When bound to MutSβ, the hairpin ends of CNG repeats resemble the stable arm and uneven bubble (Fig. 2b). The hairpin loop (turn) after the 4:1 uneven bubble has little interaction with MutSβ and consists of 3 and 6 nt at the hairpin end of (CAG)_25_ and (CTG)_25_, respectively (Fig. 3g, Extended Data Fig. 3c). As observed with the hairpin stems, the bases opposite the 4-nt insertion between YK and F42 are devoid of protein interactions and stabilized mainly by base stacking. In short, MutSβ uses the same YK motif (MSH3) and F42 (MSH2) to sandwich a four-nucleotide insertion in IDLs, uneven bubbles and hairpin ends of CNG repeats (Fig. 3a, Extended Data Fig. 5). The CCG and CGG repeats are likely bound by MutSβ similarly to (CAG)n and (CTG)n with mismatched G:G or C:C forming B-like DNA (Fig. 1c-d, Extended Data Fig. 3d) surrounding slipped-out uneven bubbles.

To find out how GAA repeats, which can form G:A Hoogsteen pairs (Extended Data Fig. 3d) like A:A but no Watson-Crick basepair at all, are still bound by MutSβ^43^, we characterized the MutSβ-(GAA)_25_ complexes by cryoEM and identified three conformational species (Conf. I - III) (Extended Data Fig. 6). In 27% of MutSβ-GAA complexes only MSH3 interacts with DNA while MSH2 is detached, and most of the DNA is disordered (Conf I). In the remaining complexes, although both MSH subunits bind DNA, 61% contain disordered DNA except for the bubble region (Conf. II), and 39% contain ordered pseudo-duplex, but the major and minor grooves are indistinct (Conf. III). These results reveal the difficulty for GAA repeats to form a stable B-like heteroduplex and explain the reduced binding affinity of MutSβ (Fig. 1c-d). The weak binding of GAA repeats by MutSβ may underlie the nonlinear relationship between MutSβ and GAA repeat expansions^1,44^.

### Tandem binding of MutSβ to large S-CNG is persistent

The unexpected MutSβ binding to the CNG hairpin stems immediately suggests that multiple MutSβ can bind a long CNG hairpin simultaneously in tandem. We hypothesize that the slip-outs resulting from increased number of CNG repeats have increased hairpin length, which in turn allows multiple MutSβ molecules to bind in tandem as observed in the EMSA analyses (Fig. 1c, Extended Data Fig. 1b). Small slip-outs are dynamic and easily reverted to homoduplexes, and a small slip-out bound by MutSβ becomes protein-free in the presence of ATP due to the MSH2 and MSH3 ATPase activities^27^. However, multiple MutSβ molecules bound to a single large hairpin are unlikely to be released simultaneously and thus would be more persistent.

To test this hypothesis, we prepared 60-repeats of CAG and CTG and found that indeed (CAG)_60_ and (CTG)_60_ (forming ∼90 bp hairpins) each can bind two, three or up to four MutSβ molecules in tandem (Fig. 4a). Furthermore, the tandem binding events were captured on negatively stained electron microscopic grids. As a control, an IDL DNA with a 3-nt insertion (Loop3) resulted in a single MutSβ bound complex (Fig. 4b, Extended Data Fig. 7a). Noticeably, 2D classifications of (CAG)_60_ and (CTG)_60_ DNA bound by three or four MutSβ are blurrier and less well ordered than these of the lower oligomeric states, which suggests that the assemblies have different registers and different basepair numbers between neighboring MutSβ molecules. Thus, the longer the repeats, the more variations of MutSβ assemblies. The heterogeneity also indicates an absence of specific MutSβ-MutSβ interactions in the tandem binding, and that oligomerization of MutSβ is mediated by the CNG repeats. This deduction is corroborated by varied arrangements of two MutSβ molecules bound to one DNA (Extended Data Fig. 2e). Based on the number of MutSβ molecules bound to (CAG)n repeats with n=18, 25, 30, 45 or 60 and structural analysis (Fig. 2a-c and 4b-c, Extended Data Fig. 7b), each MutSβ in average occupies ∼17 CNG repeats (or ∼25 bp pseudo-duplex) in the stem region and ∼9 repeats at the hairpin end. Therefore, S-CNG beyond the threshold of 40 repeats, which drives increased repeat instability^16,17^, can be bound by three or more MutSβ molecules simultaneously.

**Fig. 4.**
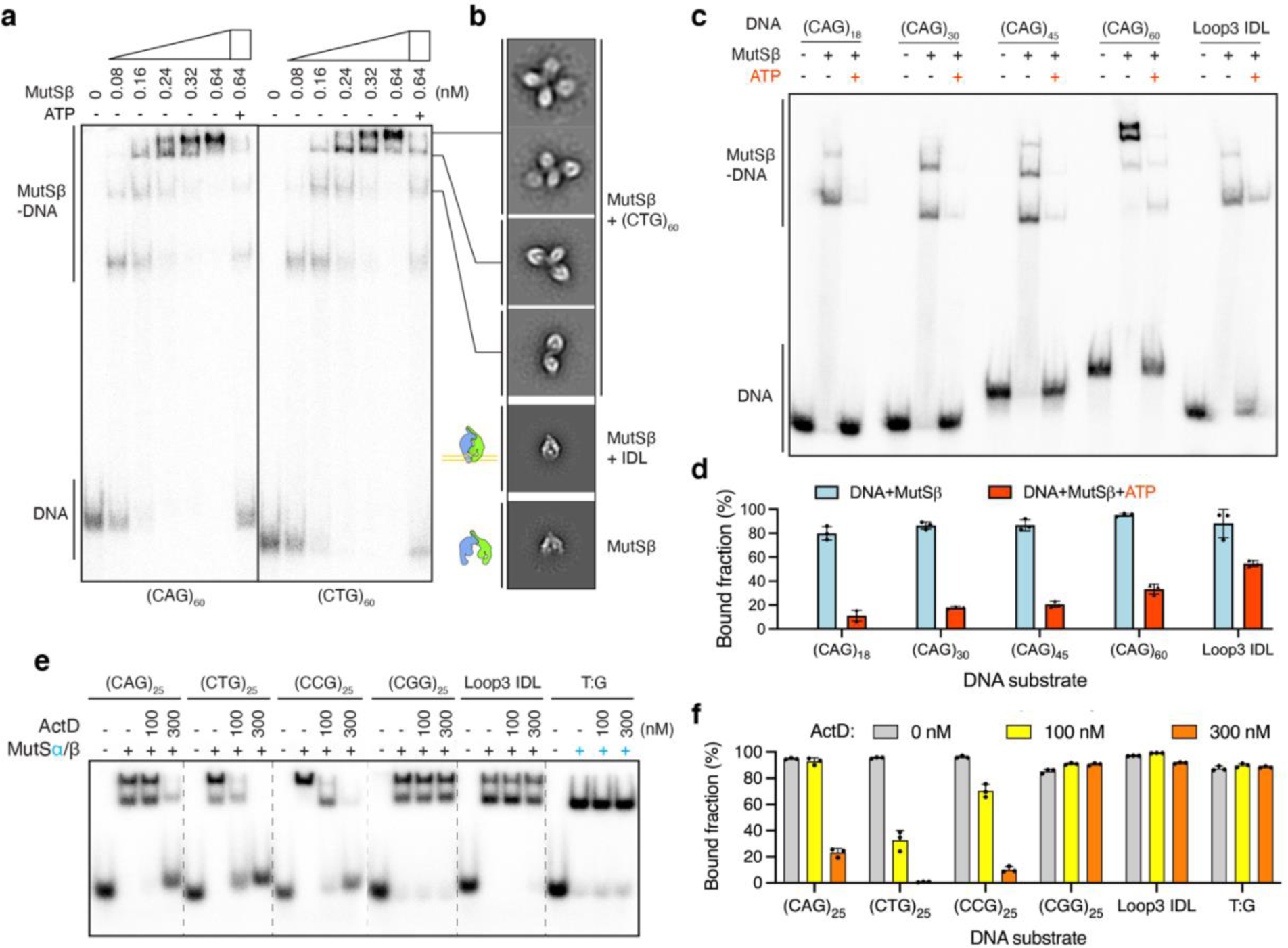
Tandem MutSβ binding to large CNG repeats and effects of ATP and ActD. **a,** EMSA analysis of tandem MutSβ binding to (CAG)_60_ and (CTG)_60_. **b,** Representative 2D projections of MutSβ, and MutSβ complexed with IDL or (CNG)_60_ from negative staining electron micrographs. **c,** EMSA assay of ATP-induced dissociation of MutSβ from different lengths of (CAG)n (n=18, 30, 45, 60) and loop3 IDL DNAs. **d,** Bar graph of the persistent MutSβ-DNA complexes detected in (**c**). **e,** ActD inhibits MutSβ binding to (CNG)_25_ without changing MutS (α or β) binding T:G or IDL mismatches. **f,** Bar graph of the persistent MutSα/β-DNA complexes detected in (**e**). In panels **d** and **f**, average of triplicate measurements is shown with the standard deviations.

As expected, ATP releases MutSβ from the pseudo-duplexes of CAG repeats as from the Loop3 IDL (Fig. 4c). However, due to asynchronous release, the longer repeats (CAG)_60_ retains more MutSβ than 18 or 30 repeats after incubation with ATP (Fig. 4d). Loop3 IDL, which contains only one MutSβ binding site, is more resistant to ATP-induced MutSβ dissociation than (CAG)_60_ bound by four MutSβ molecules (Fig. 4c-d). Tight binding of IDL ensures the efficient mismatch repair. Due to many N:N mismatches and the dynamic nature of CNG pseudo-duplex, MutSβ binding of (CNG)n hairpins is weaker than IDL, and this weaker association highlights the dependence on CNG-repeat length and tandem binding of multiple MutSβ to resist the ATP release and activate the MutLγ endonuclease, which initiates the repeat instability.

### Screening for inhibitors of MutSβ binding to CNG repeats

The unexpected finding of MutSβ binding to CNG pseudo-duplexes with every third base pair being an N:N mismatch provides a key distinction between the essential function of MutSβ in replication-associated mismatch repair and its pathological role in replication-independent repeat expansions. This division immediately suggests a simple screen for novel inhibitors that prevent the pathologic binding of MutSβ to CNG repeats but not the usual mismatches.

Previously napthyridine azaquinolone, which intercalates in CAG repeats by forming specific hydrogen bonds with A and G and flipping out basepairing partner C’s, has been reported to inhibit (CAG)n expansion^45,46^. Suspecting that DNA intercalators may prevent uneven bubble formation by stabilizing the pseudo-duplex rigidity and also cause the CNG pseudo-duplexes to deviate from the B-form conformation preferred by MutSβ, we have tested both general and N:N mismatch specific intercalators in inhibiting MutSβ binding. The commonly used DNA intercalator chloroquine (CLQ) and chromomycin A3-Ni (CMA3-Ni) reported to be a CCG specific intercalator^47^ can partially block MutSβ binding to CNG repeats at mM concentrations, but they interfere with MutS-DNA interactions in normal MMR (Extended Data Fig. 8a-b). However, actinomycin D (ActD), which prefers to intercalate at T:T mismatches^48,49^, inhibits MutSβ binding to CTG as well as CCG and CAG repeats but not to CGG or IDLs (Fig. 4e-f). At 300 nM, ActD abolishes MutSβ binding to (CTG)_25_ and (CTG)_60_ (Fig. 4e-f, Extended Data Fig. 8c). The high local concentrations of CNG repeats in expanded TNR loci may increase binding of such intercalators. We anticipate another beneficial effect of CNG-specific intercalators *in vivo* as the intercalator-stabilized pseudo-hairpins are potential targets for removal by DNA structure-specific nucleases and thus may lead to repeat contraction as observed^45,46^. Although the toxicity of ActD and brain-blood barrier prevent using it to treat repeat-expansion patients, the simple inhibitor screening promises a ready method to find effective inhibitors specific for TNR expansions.

### A model of persistent MutSβ binding leading to repeat expansion

Analogous to MMR, in which MutS binding to a mismatched DNA in the presence of ATP leads to MutL recruitment and activation of the MutL endonuclease activity^50,51^, tandem and persistent binding of MutSβ to the heteroduplexes of extruded large CNG hairpins may lead to recruitment of MutL (α or γ, and γ in particular) and DNA nicking. We suggest that MutSβ participates in CNG repeat expansion in two separate steps, repeat size-dependent DNA nicking and single-repeat expansion and contraction during error-prone repair synthesis.

Chromatin DNA is negatively supercoiled and under torsional stress when actively transcribed, which favors extrusion of inverted repeats in cruciforms^52,53^. Similarly, CNG repeats have the potential to form extruded hairpins or S-DNA in genomic DNA. In the presence of nucleosomes, S-DNA of short CNG repeats is disfavored and would quickly revert to homoduplex. Even if a short S-DNA hairpin is recognized and bound by MutSβ, the association must be transient because of the MutS ATPase activity. With long repeats, formation of a single large extrusion, however, is energetically more favorable than many small clusters of S-DNA (Fig. 5a). Moreover, a large extruded CNG hairpin can be stabilized and trapped by tandem bound MutSβ molecules in the presence of ATP (Fig. 4a-d). Formation of S-CNG repeats is potentially enhanced during transcription due to torsional stresses and the presence of single-stranded DNA^53^. Recruitment of MutL by MutSβ leads to stochastic DNA nicking by MutL (Fig. 5a).

**Fig. 5.**
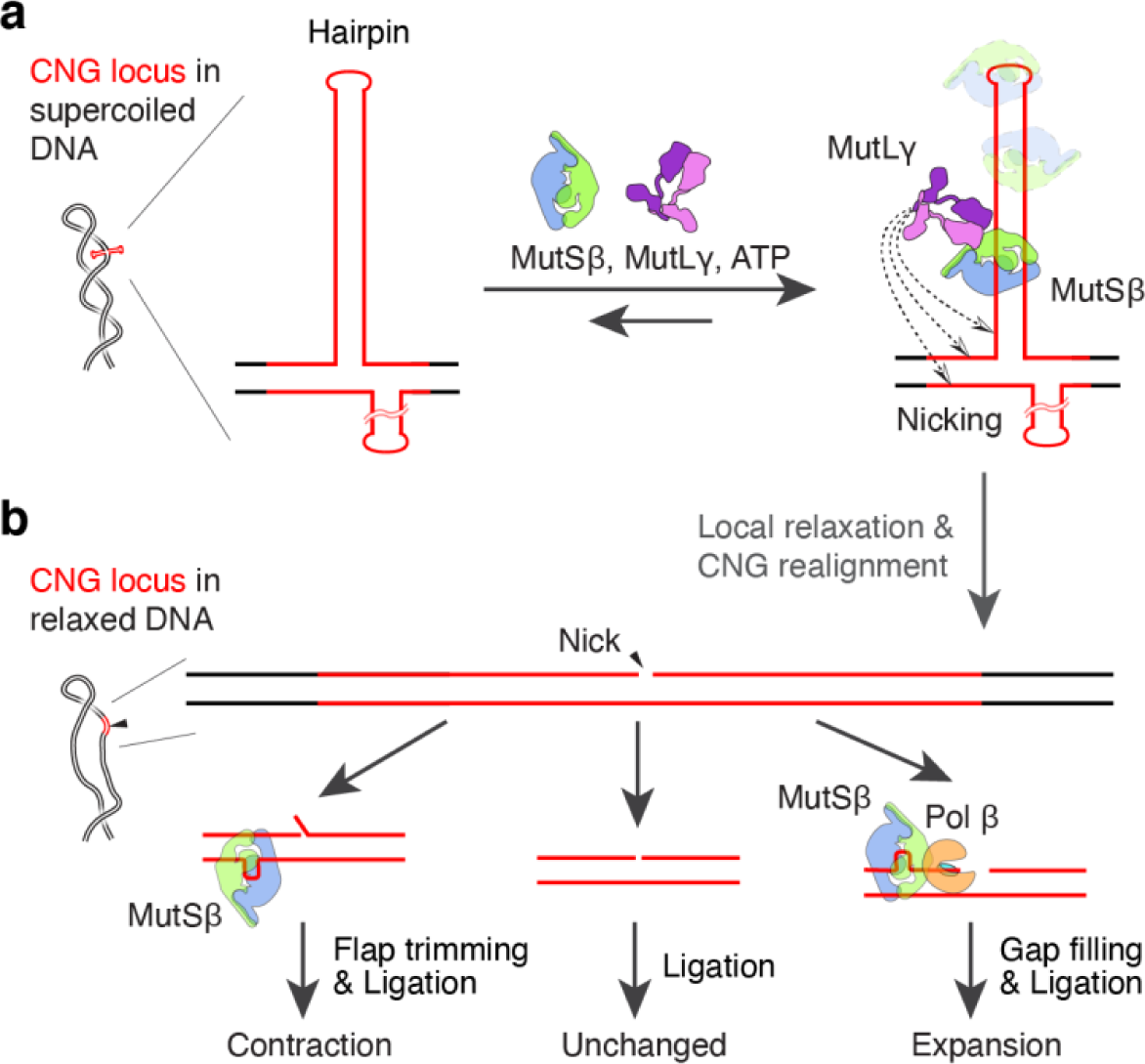
MutSβ and MutLγ dependent expansion of CNG repeats. **a,** Long extruded hairpins of (CNG)n repeats form in negatively supercoiled DNA and are stabilized by tandem binding of multiple MutS molecules. In the presence of ATP, the persistently bound MutSβ recruits and activates MutLγ endonuclease to nick the DNA (indicated by dashed arrows). **b,** Nicked DNA is relaxed, and the CNG repeats realign to form normal base-pairs. The nicked DNA may be repaired by simple re-ligation. The occurrence of a single unit slip-out is proportional to the repeat length and may lead to expansion or contraction of the repeats.

After the initial nicking, MutSβ can play a second role in CNG repeat expansion. Nicked DNA is locally relaxed, which leads to reannealing of homoduplex and release of MutSβ. As repair processes temporarily keep the nicked DNA nucleosome-free, repetitive sequences are prone to misalignment. A single repeating unit slip-out (1-us) surrounded by perfectly base paired DNA is less costly than 2-us or larger slip-outs to form. The probability of 1-us increases with the repeat length whether (CAG/CTG)n or (CCG/CGG)n repeats. Such 1-us is a typical IDL recognized and stabilized by MutSβ. If the slip-out occurs in the continuous strand, the nicked strand may be trimmed by a nuclease before re-ligation and thus result in contraction (Fig 5b). Alternatively, if the 1-us occurs in the nicked strand, interactions of MutSβ with the gap-repair DNA pol β lead to the nick-induced DNA synthesis as reported^54,55^, and expansion of one repeating unit at a time^35^ (Fig. 5b). Once a gain or loss of one repeat unit is established in one strand, another round of DNA nicking and MMR can result in expansion or contraction in both strands.

## Conclusion

Our proposal of two-step involvement of MutSβ in CNG expansion provides a testable model and suggests new research directions previously unexplored to elucidate expansion mechanisms and to search for disease treatments. The unexpected finding that MutSβ binds well to pseudo-duplexes of S-CNG DNA hairpins explains why >40 repeats of CNG increase the probability of expansion and pathogenicity. The dependence of tandem binding of multiple MutSβ molecules to long extruded CNG repeats that resist ATP release correlates well with the dependence of expansion on the high expression level of MutSβ^15^. Currently the ATPase activity of MutSβ is the prime target for drug development, but ATPase inhibitors will affect TNR expansions as well as normal MMR. Our findings suggest that there are CNG specific inhibitors like ActD, which can block TNR expansions without changing MMR.

## Supporting information

Methods,Suppmental Figs and Tables

## Acknowledgements

The authors thank Drs. R. Craigie, M. Gellert, R. Lahue, D. Leahy and K. Usdin for critical reading of the manuscript. This work utilized the Cryo-Electron Microscopy Core facility, NIDDK, and the NIH Multi-Institute Cryo-EM Facility (MICEF). This research was supported by National Institute of Diabetes, Digestive and Kidney Disease (DK036119) to W.Y.

## Author contributions

J.L. carried out biochemical and structural studies; H.W. helped with cryoEM data acquisition, J.L. and W.Y. wrote the paper, and all adhere to the “Inclusion & Ethics” regulations.

## Data and Code Availability

CryoEM maps and structure coordinates of MutSβ complexes have been deposited in the Electron Microscopy Data Bank (EMDB) and Protein Data Bank (PDB) under accession numbers EMD-42359 and 8ULL (CAG stem (1-us)), EMD-42360 and 8ULO (CAG stem (2-us)), EMD-42361 and 8ULP (CAG end), EMD-42362 and 8ULQ (CTG stem (1-us)), EMD-42367 and 8ULV (CTG stem (2-us)), EMD-42368 and 8ULW (CTG end), EMD-42369 and 8ULX ((CAG)_2_/CAG), and EMD-42370 and 8ULY ((CAG)_2_/T_3_). These data will be released immediately upon publication. Other research materials reported here are available upon request.

## Inclusion & Ethics

The authors have applied inclusion and ethics whenever possible.

