## Supplementary material for "Tandem MutSβ binding to long extruded DNA trinucleotide repeats underpins pathogenic expansions": Methods,Suppmental Figs and Tables

#### Protein expression and purification

The expression plasmids of MutSβ originate from a previous study[2] and the expression method is the same. Briefly, full-length (FL) (MSH2: 1-934 aa; MSH3: 1-1125 aa) and MSH3 N-terminus trimmed (ΔN, 211-1125 aa) MutSβ complex were expressed in Hi5 insect cells for 48h before harvest.

The cell pellet was re-suspended in buffer A (25 mM Tris-Cl pH 7.4, 1M NaCl, 5% glycerol, 1 mM Tris-(2-carboxyethyl)-phosphine (TCEP), 10 mM imidazole) and lysed by sonication. The clarified supernatant was applied to a $Ni^{2+}$ affinity column (GE Healthcare). The column was washed with 10 column volume (CV) of buffer A, then 50 CV of 90% buffer A and 10% buffer B (25 mM Tris-Cl pH 7.4, 500 mM NaCl, 5% glycerol, 1 mM TCEP, 300 mM imidazole). The target proteins were eluted with 100% buffer B. The fractions containing target proteins were pooled and mixed with PreScission protease (protease : protein mass ratio 1:50) for 3h at 4℃ to remove the $His_8$-MBP tag. The protein sample was diluted to 100 mM NaCl by dilution buffer (25 mM Tris-Cl pH 7.4, 10% glycerol, 1 mM TCEP, 0.5 mM EDTA) and further purified over a Heparin column (GE Healthcare), with a linear NaCl gradient from 100−1000 mM plus 25 mM Tris-Cl pH 7.4, 5% glycerol, 0.5 mM EDTA, 1 mM TCEP. Purified MutSβ proteins were concentrated to ~2 mg/ml, mixed with glycerol to 50% final concentration, aliquoted and stored at -80℃.

A similar cloning strategy of MutSβ[2] was used for the MutS$\alpha$ complex. FL MSH2 (1-934 aa) and N-terminal truncated and $His_8$-MBP-PreScission tagged MSH6 (342-1360 aa) were cloned into pFastbac Dual vector under the p10 and pPH promoters, respectively. Due to the unequal expression level of the two subunits, another plasmid expressing FL MSH2 with an N-terminal Glutathione S-transferase (GST) tag followed by the PreScission protease site in pFastbac was made. They were co-expressed in Hi5 insect cells for 48h.

The cell pellet was re-suspended in buffer C (25 mM Tris-Cl pH 7.4, 1M NaCl, 5% glycerol, 1 mM TCEP, 30 mM imidazole) and lysed by sonication. The clarified supernatant was applied to a $Ni^{2+}$ affinity column (GE Healthcare). The column was stepwise washed with 0%, 10%, 30% buffer B (25 mM Tris-Cl pH 7.4, 500 mM NaCl, 5% glycerol, 1 mM TCEP, 300 mM imidazole) mixing with buffer C, each for 30 CV. The target proteins were then eluted with 100% buffer B. The fractions containing target proteins were pooled and mixed with PreScission protease

(protease : protein mass ratio 1:50) for 3h at 4℃ to remove the $His_8$-MBP and GST tags. The protein sample was diluted to 100 mM NaCl by dilution buffer (25 mM Tris-Cl pH 7.4, 10% glycerol, 1 mM TCEP, 0.5 mM EDTA) and further purified over a monoQ column (Cytiva) with the same buffer as for Hairpin purification of MutSβ. The target proteins were further loaded into superdex-200 (Cytiva) pre-equilibrated with buffer (25 mM HEPES pH 7.5, 100 mM KCl, 0.1 mM EDTA, 5 mM $MgCl_2$, 5% glycerol, 1mM TCEP). Purified MutS$\alpha$ proteins were concentrated to ~9 mg/ml, mixed with glycerol to 50% final concentration, aliquoted and stored at -80℃.

**DNA substrate preparation**

All DNA oligonucleotides (Extended Data Table 2) were purchased from IDT (http://www.idtdna.com) and further purified by 8%-15% Urea-PAGE and desalted (BioRad). The DNA substrates were annealed in buffer (20 mM HEPES pH 7.5, 100 mM KCl).

**Cryo-EM grid preparation**

$(CAG)_{25}$ was mixed with MutSβ (ΔN), and all the other substrates ($(CTG)_{25}$, d40h30, TriArm_9-6, $(GAA)_{25}$, $(CAG)_2$/CAG, $(CAG)_2/T_3$) were mixed with FL MutSβ. The protein: DNA molar ratio is 2:1 for d40h30, and 1:1.2 for all the other DNAs. The mixture was incubated on ice for 20 minutes and then changed (by dilution and concentration by centrifugation or dialysis) into the grid freezing buffer (20 mM HEPES pH 7.5, 100 mM KCl, 5 mM $MgCl_2$, 1 mM DTT). The final concentration of the sample was adjusted to 2.8 mg/ml for MutSβ (ΔN) and 0.3 mg/ml for FL MutSβ, and 0.02-0.05% octyl-beta-glucoside ($\beta$-OG) was included. Grids were prepared using a Vitrobot (Thermo Fisher Scientific) at 4℃ and 100% humidity. 2.5 $\mu$l sample was loaded onto Quantifoil R 1.2/1.3 Au 300 mesh grids (EMS) pre-cleaned with a PELCO easiGlow glow discharge system (PELCO). The blotting force was 4 and the time was 2.5 s.

**Cryo-EM data collection**

Data of MutSβ complexed with $(CAG)_{25}$, $(CTG)_{25}$, d40h30, and TriArm_9-6 were collected on the MICEF (Multi-Institute Cryo-EM Facility at NIH) Titan Krios transmission electron microscope (Thermo Fisher Scientific) with a K3 Summit direct electron detector (Gatan) in the super-resolution mode (105k magnification, pixel size = 0.4165 Å). The exposure rate ranged from 27.7 to 31.2 $e^-$/ $Å^2$/s, with a total exposure of 45.76 to 56.28 $e^-$/ $Å^2$.

Data of MutSβ complexed with $(GAA)_{25}$ and bubble DNAs ($(CAG)_2$/CAG and $(CAG)_2/T_3$) were collected on the NIDDK Glacios electron microscope (Thermo Fisher Scientific) with a K3

Summit direct electron detector in the super-resolution mode (45k magnification, pixel size = 0.434 Å). The exposure rate was 28.3 $e^-$/ Å$^2$/s, with a total exposure of 70.0 $e^-$/ Å$^2$.

All data sets were collected using SerialEM[56]. The defocus range was set to -0.5 to -1.5 $\mu$m (see Extended Data Table 1 for more details).

**Cryo-EM data processing**

Nearly all procedures were carried out in RELION (version 3.1.4)[57,58]. The raw super-resolution movies were binned by 2-fold to pixel size of 0.868 Å. and beam-induced motion was corrected by MotionCor2 (version 1.3.0)[59] in 7 × 7 patches. The defocus values were estimated with the non-dose-weighted micrographs by Gctf (version 1.06)[60]. Micrographs with estimated resolution worse than 3.5 Å were discarded. Gautomatch (http://www.mrc-lmb.cam.ac.uk/kzhang/Gautomatch) was used to pick particles with the template generated from the crystal structure of MutSβ (PDB ID: 3THX). A box size of 160 pixels was used to extract 2×-binned particles (pixel size = 1.736 Å) from the dose-weighted micrographs. After two rounds of 2D classification (mask diameter = 220 Å), selected particles were subjected to 3D classification. Particles in good classes were re-extracted without binning (box size = 320 pixels. pixel size = 0.833 Å and 0.868 Å for Krios datasets and Glacios datasets, respectively). Particles in similar classes were combined for auto-refinement, followed with CTF refinement and Bayesian polishing to improve the resolution (Extended Data Fig. 2, 4).

To separate different DNA-binding modes and improve map quality, 3D classification without alignment was applied to the DNA and surrounding protein domains. Several different Tau2Fudge factor values and different class numbers were tested in parallel, and resulting good classes were kept for further refinement. Several rounds of local classification were performed in some cases to separate different conformational species and get the best maps. All the reported resolutions are of the overall maps and follow the criterion of gold-standard Fourier shell correlation (FSC)=0.143. Local resolution distribution maps were generated by RELION.

Data processing workflows for (CNG)n, uneven bubble DNAs, and (GAA)n are summarized in Extended Data Fig. 2-3 and 6, respectively. The d40h30 DNA with $(CAG)_{30}$ in the middle and $(CAG)_{25}$ DNA datasets were separately processed initially. Since the $(CAG)_n$ hairpins bound by MutSβ shared the same conformation, they were combined later as shown in Extended Data Fig. 2a. To determine the populations of MutSβ molecular bound at the hairpin stem or hairpin end, we used the d40h30 dataset only and counted particles in the CAG stem-binding and CAG end-binding classes separately.

**Model building and refinement**

Refined cryo-EM maps were treated with DeepEMhancer[61] for model building, and the unsharpened raw RELION maps were used for refinement and structure validation.

The crystal structure of MutSβ complexed with Loop3 IDL (PDB ID: 3THX) was used as a template for model building. Once the CAG stem (1-us) model was built, it was used as the initial model for others. DNAs were manually built in Coot (version 0.8.9.2 EL)[62]. The structures were then improved with Real-Space Refinement[63] in PHENIX (version 1.20.1-4487-000)[64]. As the DNAs contain many mismatched pairs, additional geometry restraints, including bond length, angles and planarity were applied during Real-Space Refinement. Because stereochemical parameters of the T:T base pairing (Saenger NO. 16) are absent in Phenix, we used the restraints derived from the 1.59 Å structure of 6BOW (PDB)[65]. The final models were obtained after iterative rounds of model adjustment and refinement using Coot and PHENIX (Extended Data Table 1). We have noted that MSH2 in MutSβ (ΔN) contains an ADP, which remained bound throughout the protein purification and cryoEM grid preparation, which agrees with a previous observation[2]. Interestingly, MutSβ (FL) is devoid of any ADP (Extended Data Fig. 7c).

All structure figures were prepared with UCSF Chimera[66] and ChimeraX[67].

**DNA structure analysis**

Coordinates of different DNAs complexed with MutSβ were analyzed using the online 3DNA 2.0 (http://web.x3dna.org/analyze_file/parameter). The helical axis of a duplex arm was the sum of all dinucleotide vectors within it. The angle between each pair of stable and mobile arms was calculated using the software https://www.omnicalculator.com/math/angle-between-two-vectors.

**Negative-staining EM analysis**

MutSβ was mixed with $(CAG)_{60}$, $(CTG)_{60}$, or Loop3 IDL at a molar ratio of 4:1, 4:1 and 1:1, respectively, in incubation buffer (20 mM HEPES pH 7.5, 100 mM KCl, 5% glycerol, 1 mM DTT) and incubated on ice for 20 min. Concentrations of MutSβ alone and MutSβ-DNA were adjusted to 0.02 mg/ml with incubation buffer. 2 $\mu$l sample solution was loaded onto a 400 mesh Cu grid with continuous carbon film (EMS) pre-cleaned with a PELCO easiGlow glow discharge system (PELCO) and incubated for 1 min at room temperature (RT). The excess solution was removed with filter paper. The grid was washed with 4 $\mu$l of 2% uranyl formate staining solution twice, stained for 1 min, and air dried at RT.

Negative-staining EM datasets were collected on a Tecnai TF20 transmission electron microscope (Thermo Fisher Scientific) using serialEM, with pixel size of 1.26 Å. 80, 422 and 478 micrographs were collected for apo MutSβ, MutSβ-Loop3 IDL and MutSβ-$(CTG)_{60}$ complexes, respectively. The micrographs were processed using RELION (version 3.1). Particles in Apo MutSβ and MutSβ-Loop3 IDL datasets were picked using Gautomatch without template and with a diameter of 200 Å. The particles on MutSβ-$(CTG)_{60}$ micrographs were of different oligomeric states and they were picked manually with a diameter of 500 Å. 13,021, 108,266, and 11,436 particles were picked from the apo MutSβ, MutSβ-Loop3 IDL and MutSβ-$(CTG)_{60}$ micrographs, respectively. Particles were extracted with 2× binning factor (pixel size = 2.52 Å) and box size = 240 pixel. For 2D classification (100 classes and T = 2), the mask diameter was set to 500 Å for the MutSβ-$(CTG)_{60}$ dataset and 360 Å for the other two.

**Electrophoretic mobility shift assay (EMSA) and dissociation constant ($K_d$) measurement**

100 nM DNA oligos were 5′-end labeled using $\gamma$-$^{32}$P-ATP (10 $\mu$Ci/$\mu$l. PerkinElmer) and T4 polynucleotide kinase (NEB) in 20 $\mu$l buffer and incubated at 37℃ for 0.5 h. The oligos were purified by ProbeQuant G-50 micro columns (Cytiva). For DNAs formed with multiple strands, the unlabeled strands were added at a molar ratio of 1.5× of the labeled one, then annealed in the annealing buffer (20 mM HEPES pH 7.5, 100 mM KCl) at a final concentration of 50 nM. The DNAs were stored at -20℃.

For $K_d$ measurement (Fig.1 and Extended Data Fig. 4b), each 40 $\mu$l reaction mix contained 10 pM $^{32}$P- labeled DNA and MutSβ (ΔN) proteins in various concentrations as indicated (figures and figure legends) in 1× EMSA buffer (25 mM HEPES pH 8.0, 100 mM NaCl, 5 mM $MgCl_2$, 0.1 mM EDTA, 20% glycerol, 2 mM TCEP, 0.2 mg/ml BSA). After incubation for 20 min at RT, 8 $\mu$l of each sample was resolved on 5% or 8% native polyacrylamide gel electrophoresis (PAGE) (29:1) at 100 V and 4℃ for 1.5 h. The radioactive bands were detected by a Typhoon PhosphorImager and quantified using ImageQuant NL (GE Healthcare). Each assay was repeated three or more times. The Junction binding curve in Fig. 1b was produced based on the EMSA of TriArm_9-9 DNA as shown in Extended Data Fig. 1d. Binding curves were fitted and Kd determined using GraphPad Prism program version 9 (GraphPad Software). Each curve and error bars were generated based on three independent measurements.

**ATP-dependent dissociation of MutSβ from IDL and (CNG)n**

The DNA oligos were $^{32}$P-labeled and EMSA were performed as for $K_d$ measurement. Medium-sized (20 cm × 20 cm) 5% (29:1) native PAGE gels were used here to better separate multiple shifted bands. For EMSA gels shown in Fig. 4a, 20 pM $^{32}$P-labeled DNA and MutSβ (ΔN) (80 to 640 pM) as indicated were mixed. After incubation at RT for 10 min, 8 $\mu$l sample was loaded into each well and electrophoresed in 1× TBE buffer at 250 V and 4℃ for 3h. For EMSA gels shown in Fig. 4c and Extended Data Fig. 7b, 100 pM $^{32}$P-labeled DNA and 600 pM MutSβ (ΔN) were mixed and incubated at RT for 10 min, and then after addition of 0.5 mM ATP incubated for another 5 min at 4℃ before loading onto the gel. The electrophoresis was done in 1× TBE buffer at 4℃ and 250 V for 2h for samples in Fig. 4c. For samples in Extended Data Fig. 7b, the running time varied according to the substrate length so that they can be clearly observed in the same gel. It was 1h, 2h, and 3h for $(CAG)_8$, $(CAG)_{25}$, $(CAG)_{60}$, respectively.

The gels were dried for 3h at 80℃ before scanning with a Typhoon PhosphorImager. The bands were quantified using ImageQuant NL (GE Healthcare).

**Intercalator inhibitory assay**

Chromomycin A3 (CMA3) (sigma, NO. C2659) and $NiSO_4$ (sigma, NO.227676) were dissolved in dimethyl sulfoxide (AmericanBio, NO. AB03091) and deionized water, respectively. Mixing CMA3 and $NiSO_4$ with 2:1 molar ratio to prepare CMA3-Ni stock solution in 20 mM. Actinomycin D (sigma, NO. A1410) was dissolved in acetonitrile (sigma, NO. 900667) to 8 mM stock solution. Chloroquine (sigma, NO. C6628) was dissolved in deionized water to 100 mM stock solution.

For DNA specificity assay in Fig. 4e and Extended Data Fig. 8a, b, 2 nM $^{32}$P-labeled $(CNG)_{25}$ DNAs or similar length of TG mismatch DNA or loop3 IDL DNA were incubated with 4 nM MutSβ proteins in 1× EMSA buffer, with or without chemicals at the indicated concentrations. After incubation for 20 min at RT, 8 $\mu$l of each sample was resolved on 5% native PAGE (29:1) at 120 V and 4℃ for 45 min. For DNA specificity assay in Extended Data Fig. 8c, 20 $\mu$l reaction mix contained 0.5 nM $^{32}$P-labeled $(CNG)_{60}$ DNAs or similar length of TG mismatch DNA or loop3 IDL DNA and 2 nM MutSβ proteins in 1× EMSA buffer, with indicated concentration of ActD.

The sample was resolved on 5% native PAGE (29:1) Medium-sized (20 cm × 20 cm) gels at 250V for 3h at 4℃.

The radioactive bands were detected by a Typhoon PhosphorImager and quantified using ImageQuant NL (GE Healthcare).

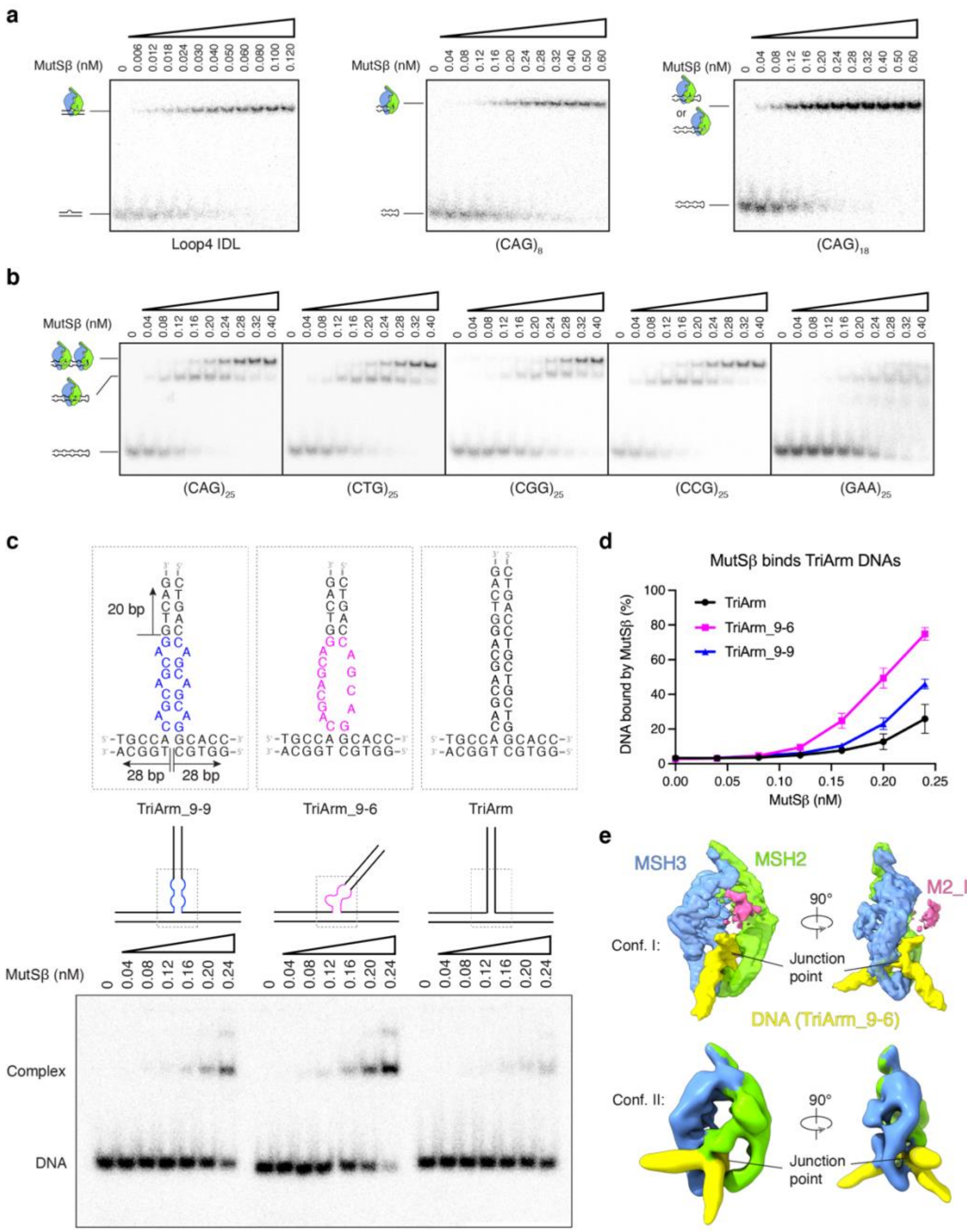


**Extended Data Fig. 1. Binding of MutSβ to S-DNAs**

**a,** EMSA gels of MutSβ binding to Loop4 IDL, $(CAG)_8$ and $(CAG)_{18}$ DNA hairpins. **b,** EMSA gels of MutSβ binding to $(CNG)_{25}$ (N=A, T, G, and C) and $(GAA)_{25}$ DNA hairpins. **c,** DNA design and EMSA gels of MutSβ binding to DNA three-way junctions. DNA sequences in dashed boxes are shown above the diagrams. **d,** Binding curves derived from the EMSA gels (**c**). Each curve shows averaged values of triplicated EMSA measurements. Error bars represent the standard deviations. **e,** Two representative conformations of the MutSβ-TriArm_9-6 complexes resulting from cryo-EM analysis. MSH2, MSH3 and DNA are colored green, blue and yellow, respectively. In Conf. I, domain II of MSH2 (M2_II, highlighted in pink) is rather flexible and partially disordered, while M2_I was not observed.

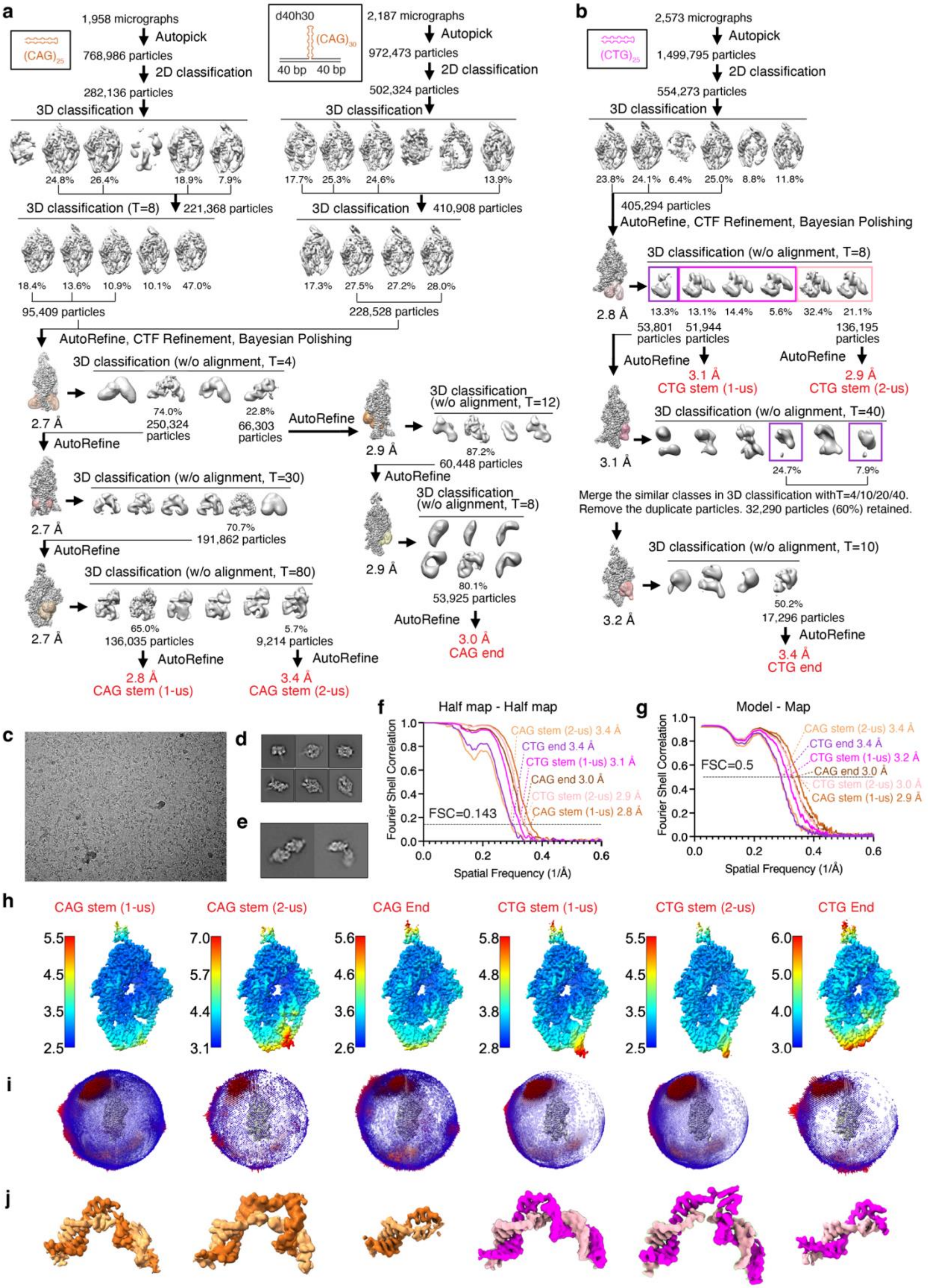

a
(CAG)25
1,958 micrographs
Autopick
768,986 particles
2D classification
282,136 particles
3D classification
24.8% 26.4% 18.9% 7.9%
3D classification (T=8)
221,368 particles
18.4% 13.6% 10.9% 10.1% 47.0%
95,409 particles
d40h30
(CAG)30
40 bp 40 bp
2,187 micrographs
Autopick
972,473 particles
2D classification
502,324 particles
3D classification
17.7% 25.3% 24.6% 13.9%
3D classification
410,908 particles
17.3% 27.5% 27.2% 28.0%
228,528 particles
AutoRefine, CTF Refinement, Bayesian Polishing
3D classification (w/o alignment, T=4)
74.0%
250,324 particles
22.8%
66,303 particles
2.7 Å
AutoRefine
3D classification (w/o alignment, T=30)
70.7%
191,862 particles
2.7 Å
AutoRefine
3D classification (w/o alignment, T=80)
65.0%
136,035 particles
5.7%
9,214 particles
2.7 Å
AutoRefine
2.8 Å
CAG stem (1-us)
AutoRefine
3.4 Å
CAG stem (2-us)
AutoRefine
2.9 Å
3D classification (w/o alignment, T=12)
87.2%
60,448 particles
AutoRefine
3D classification (w/o alignment, T=8)
2.9 Å
80.1%
53,925 particles
AutoRefine
3.0 Å
CAG end
b
(CTG)25
2,573 micrographs
Autopick
1,499,795 particles
2D classification
554,273 particles
3D classification
23.8% 24.1% 6.4% 25.0% 8.8% 11.8%
405,294 particles
AutoRefine, CTF Refinement, Bayesian Polishing
3D classification (w/o alignment, T=8)
2.8 Å
13.3% 13.1% 14.4% 5.6% 32.4% 21.1%
53,801 particles
51,944 particles
136,195 particles
AutoRefine
AutoRefine
3.1 Å
CTG stem (1-us)
2.9 Å
CTG stem (2-us)
3D classification (w/o alignment, T=40)
3.1 Å
24.7% 7.9%
Merge the similar classes in 3D classification withT=4/10/20/40.
Remove the duplicate particles. 32,290 particles (60%) retained.
3D classification (w/o alignment, T=10)
3.2 Å
50.2%
17,296 particles
AutoRefine
3.4 Å
CTG end
c
d
e
f
Half map - Half map
Fourier Shell Correlation
Spatial Frequency (1/Å)
FSC=0.143
CAG stem (2-us) 3.4 Å
CTG end 3.4 Å
CTG stem (1-us) 3.1 Å
CAG end 3.0 Å
CTG stem (2-us) 2.9 Å
CAG stem (1-us) 2.8 Å
g
Model - Map
Fourier Shell Correlation
Spatial Frequency (1/Å)
FSC=0.5
CAG stem (2-us) 3.4 Å
CTG end 3.4 Å
CTG stem (1-us) 3.2 Å
CAG end 3.0 Å
CTG stem (2-us) 3.0 Å
CAG stem (1-us) 2.9 Å
h
CAG stem (1-us)
CAG stem (2-us)
CAG End
CTG stem (1-us)
CTG stem (2-us)
CTG End
5.5 4.5 3.5 2.5
7.0 5.7 4.4 3.1
5.6 4.6 3.6 2.6
5.8 4.8 3.8 2.8
5.5 4.5 3.5 2.5
6.0 5.0 4.0 3.0
i
j

**Extended Data Fig. 2. cryoEM data processing of MutSβ-(CAG)n and $(CTG)_{25}$ complexes**

**a-b,** Data processing flowcharts of MutSβ complexed with $(CAG)_{25}$, $(CAG)_{30}$ (d40h30) (**a**) or $(CTG)_{25}$ (**b**). The maps labeled in red were used for model building and refinement. The classes boxed in magenta, salmon and purple represented 1-us and 2-us CTG stem binding, and CTG end binding, respectively. **c,** A representative micrograph. **d** and **e,** Results of reference-free 2D class averaging of MutSβ-$(CAG)_{25}$ with a mask diameter of 220 Å (**d**) and tandem binding of two MutSβ to each d40h30 DNA with a mask diameter of 360 Å (**e**). **f-g,** FSC analysis of the map quality and resolution (**f**) and model-map fit of each complex (**g**). **h-j**, For each MutSβ-DNA complex, a surface presentation of the map local resolution with the scale bar on the side (**h**), angular distributions of particles used for the final three-dimensional reconstruction (**i**), and local map of DNA (**j**) are shown.

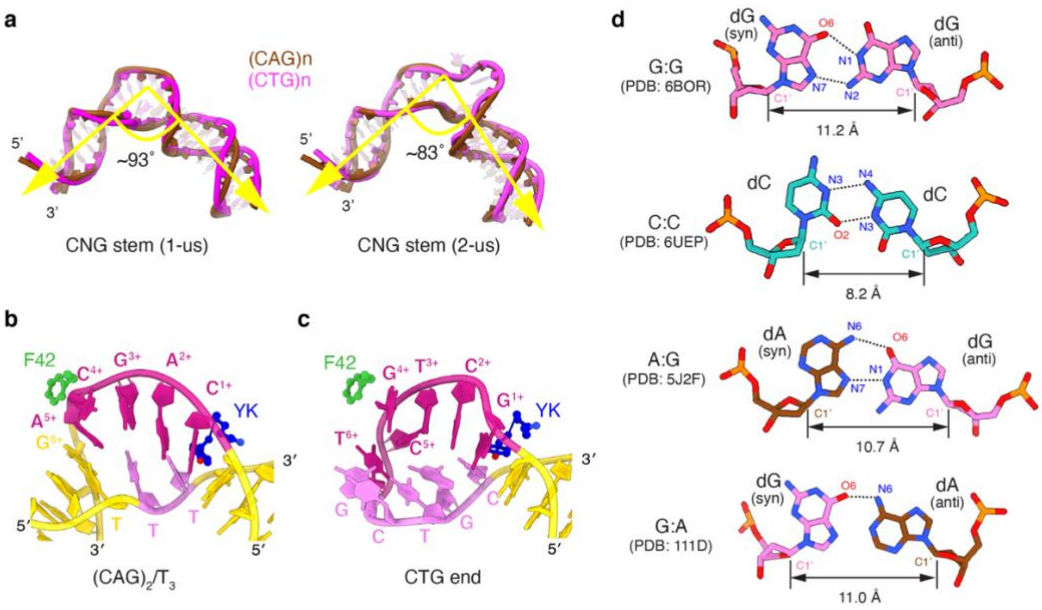


**Extended Data Fig. 3. DNA structures in MutSβ-$(CNG)_{25}$ complexes**

**a,** Different DNA bending angles of 1-us and 2-us bound to MutSβ. The angles between stable and mobile arms were determined using web 3DNA 2.0. **b,** The uneven bubble structure of $(CAG)_2/T_3$ DNA. **c,** The hairpin loop structure of $(CTG)_{25}$. The coloring scheme is the same as Fig. 3a-f. **d,** Structures of potential mismatched pairs in (CCG)n, (CGG)n and (GAA)n pseudoduplexes. They were extracted from existing crystal structures (PDB: 6BOR, 1.84 Å; PDB: 6UEP, 2.05 Å; PDB: 5J2F, 2.10 Å; PDB: 111D, 2.25 Å;).

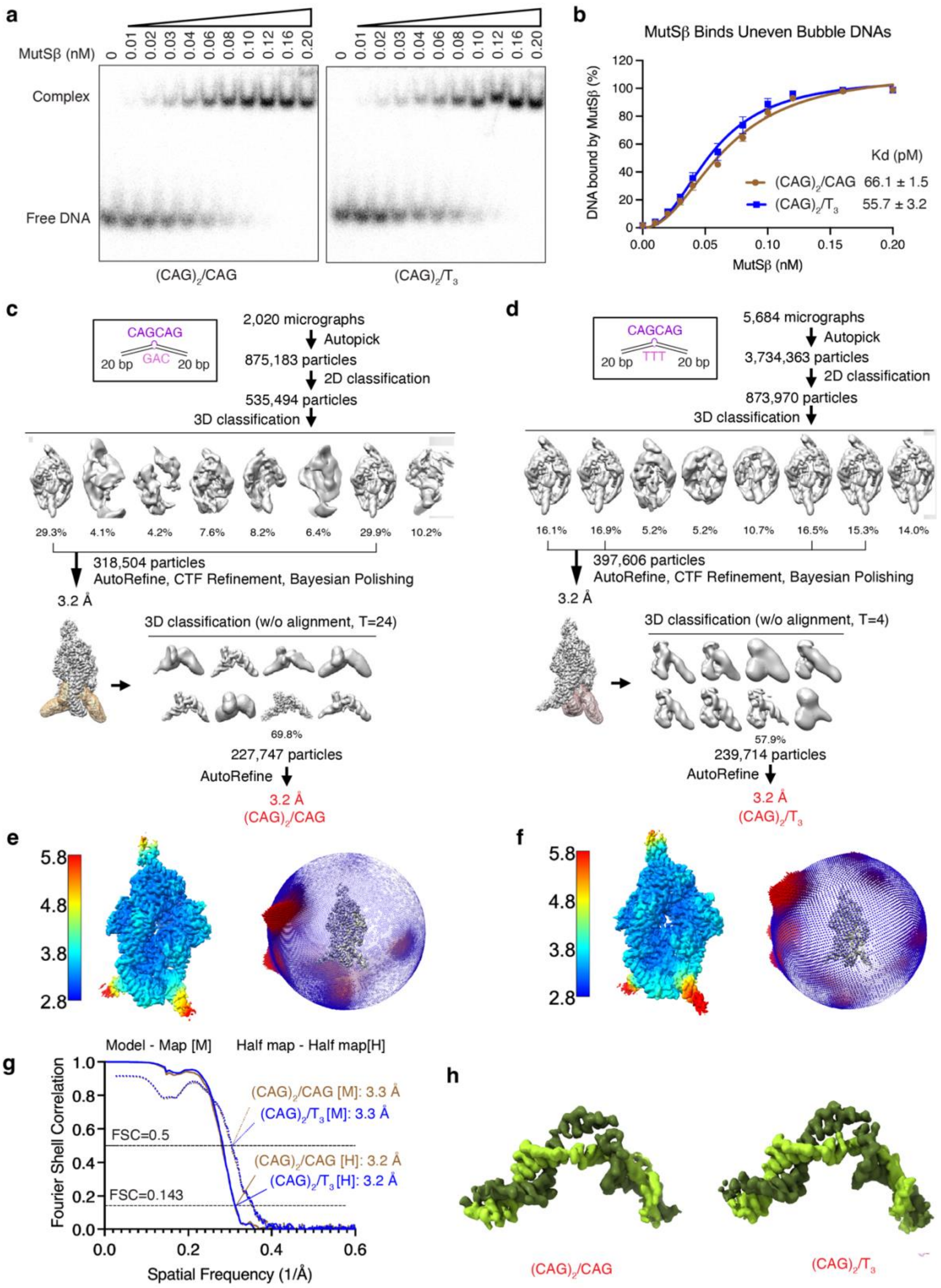
a
MutSβ (nM)
0 0.01 0.02 0.03 0.04 0.06 0.08 0.10 0.12 0.16 0.20
Complex
Free DNA
(CAG)2/CAG
(CAG)2/T3
b
MutSβ Binds Uneven Bubble DNAs
DNA bound by MutSβ (%)
MutSβ (nM)
Kd (pM)
(CAG)2/CAG 66.1 ± 1.5
(CAG)2/T3 55.7 ± 3.2
c
CAGCAG
GAC
20 bp
20 bp
2,020 micrographs
Autopick
875,183 particles
2D classification
535,494 particles
3D classification
29.3% 4.1% 4.2% 7.6% 8.2% 6.4% 29.9% 10.2%
318,504 particles
AutoRefine, CTF Refinement, Bayesian Polishing
3.2 Å
3D classification (w/o alignment, T=24)
69.8%
227,747 particles
AutoRefine
3.2 Å
(CAG)2/CAG
d
CAGCAG
TTT
20 bp
20 bp
5,684 micrographs
Autopick
3,734,363 particles
2D classification
873,970 particles
3D classification
16.1% 16.9% 5.2% 5.2% 10.7% 16.5% 15.3% 14.0%
397,606 particles
AutoRefine, CTF Refinement, Bayesian Polishing
3.2 Å
3D classification (w/o alignment, T=4)
57.9%
239,714 particles
AutoRefine
3.2 Å
(CAG)2/T3
e
5.8
4.8
3.8
2.8
f
5.8
4.8
3.8
2.8
g
Model - Map [M]
Half map - Half map[H]
Fourier Shell Correlation
Spatial Frequency (1/Å)
FSC=0.5
FSC=0.143
(CAG)2/CAG [M]: 3.3 Å
(CAG)2/T3 [M]: 3.3 Å
(CAG)2/CAG [H]: 3.2 Å
(CAG)2/T3 [H]: 3.2 Å
h
(CAG)2/CAG
(CAG)2/T3

**Extended Data Fig. 4. Binding of uneven bubbles by MutSβ and cryoEM data processing**

**a,** EMSA gels of MutSβ binding to uneven bubble DNAs. **b,** Binding curves derived from EMSA analysis (**a**). Each data point and error bar resulted from three independent measurements. **c-d,** Data-processing flowcharts of $(CAG)_2$/CAG (**c**) and $(CAG)_2/T_3$ (**d**) cryoEM datasets. The maps labeled in red were used for the model building and refinement. **e-f,** Surface presentation of the map local resolution with the scale bar on the side and angular distributions of particles used for the final three-dimensional reconstruction of $(CAG)_2$/CAG (**e**) and $(CAG)_2/T_3$ complexes (**f**). **g,** FSC analysis of the map quality and resolution (solid lines) and the model-map fit (dash lines). **h,** Local map of uneven bubble DNAs bound by MutSβ.

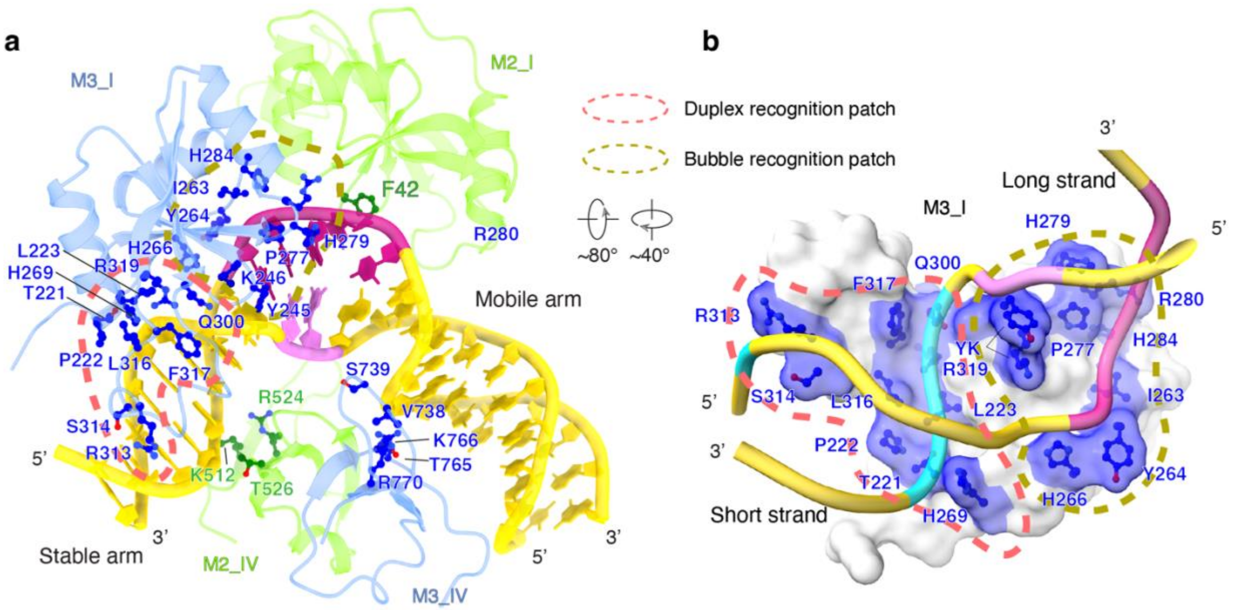


**Extended Data Fig. 5. Protein-DNA interaction of MutSβ-DNA complex**

**a,** Protein-DNA interactions in MutSβ-(CAG)n (1-us) complex. The structures are colored in the same scheme as in Fig.3. The key residues for DNA binding are shown as sticks. The residues on M3_I can be grouped into two clusters, duplex recognition patch (orange dashed line) and bubble recognition patch (olive dashed line). **b,** The DNA binding surface in M3_I. M3_I is shown as a semi-transparent light grey surface with the DNA binding residues highlighted in blue. For simplicity, only DNA backbones are shown. The nucleotides interacting with M3_I are highlighted in cyan. Two interfaces are marked with dashed circles (pink and olive).

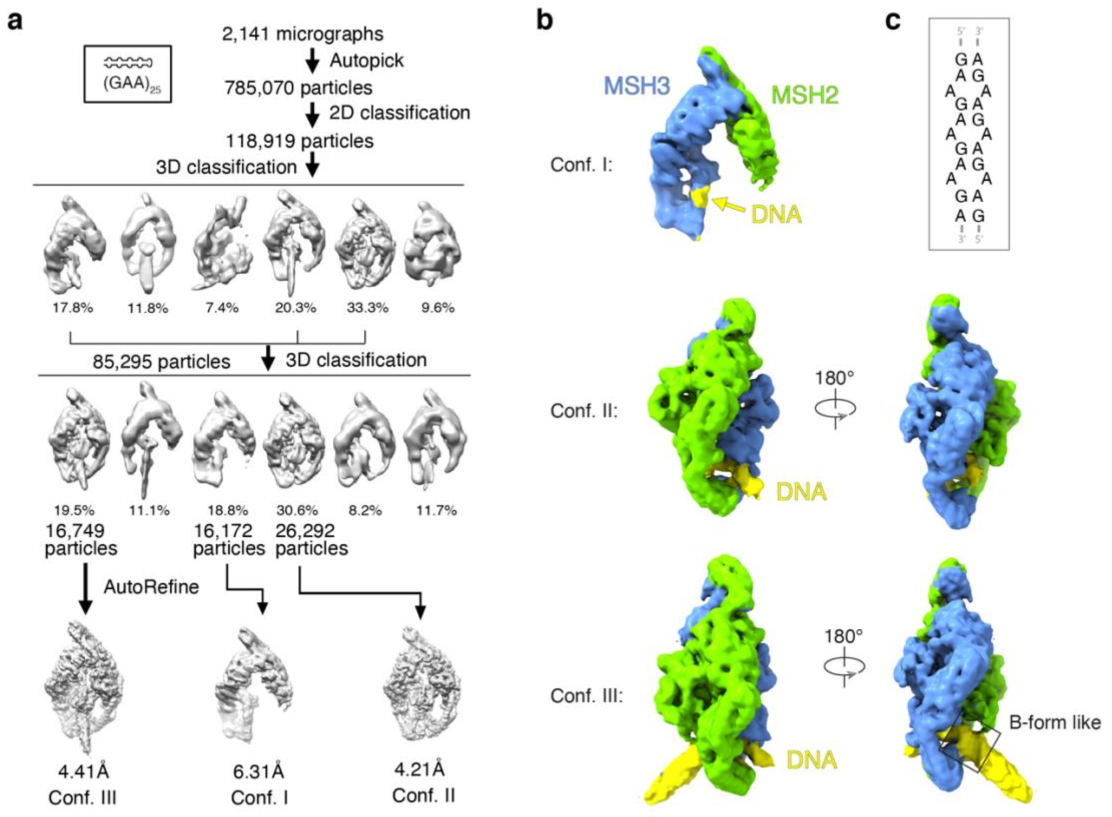


**Extended Data Fig. 6. Data processing of the MutSβ-(GAA)25 complex**

**a,** Data processing flowchart of the MutSβ-(GAA)25 complex. **b,** Three conformations were isolated from the dataset. Corresponding cryoEM maps are shown with MSH2, MSH3 and DNA colored green, blue and yellow, respectively. **c,** Supposed duplex formed by GAA repeats. G:A may form pairing as shown in Extended Data Fig. 3d, A:A may form pairing as shown in Fig. 2e.

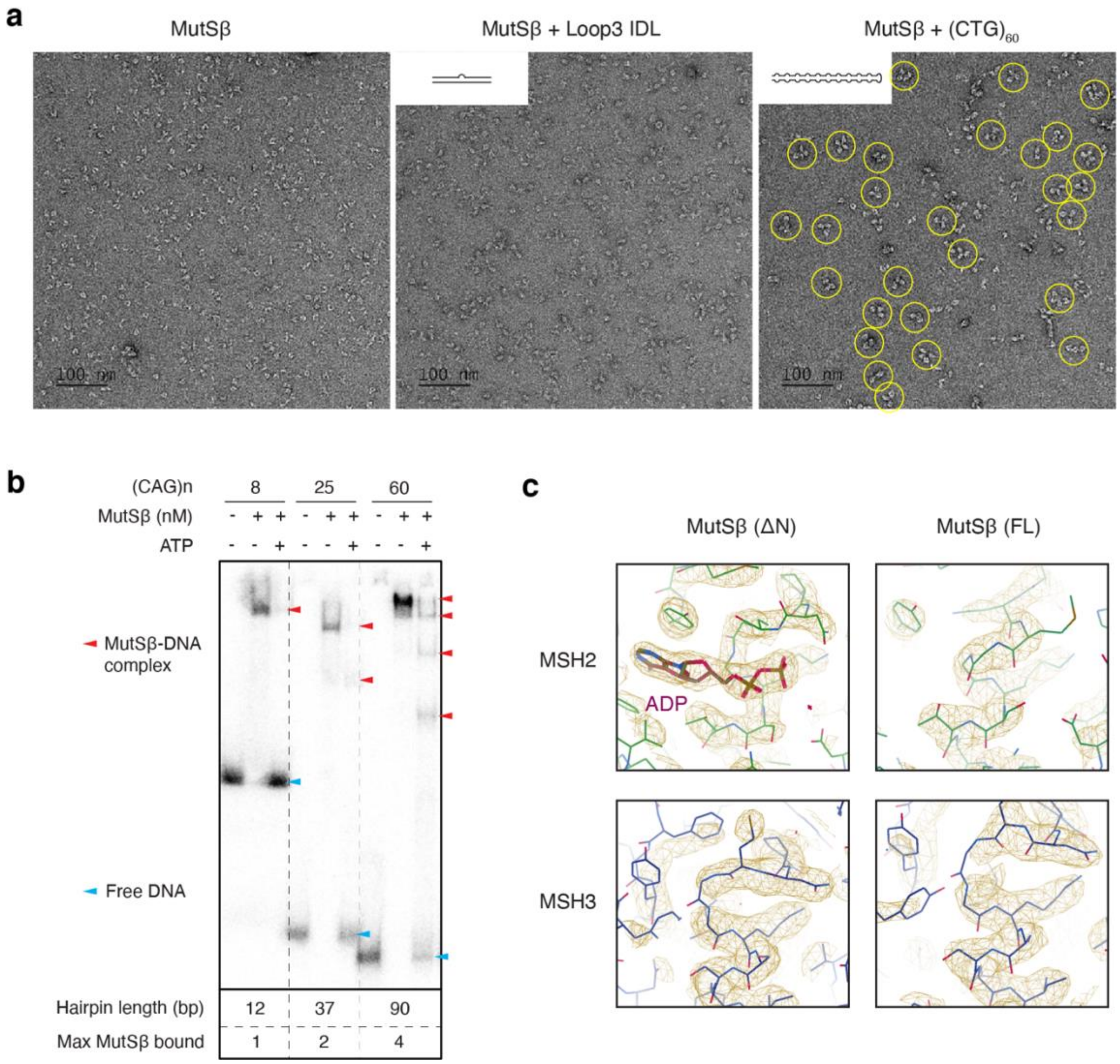


**Extended Data Fig. 7. Tandem binding of MutSβ to long (CNG)n hairpins**

**a,** Representative negatively stained micrographs of apo MutSβ alone, MutSβ complexed with Loop3 IDL or $(CTG)_{60}$. Particles containing multiple MutSβ on each $(CTG)_{60}$ are circled in yellow. **b,** EMSA gels of MutSβ binding to different (CAG)n repeats (n= 8, 25, 60). Free DNAs and MutSβ-DNA complexes are indicated by blue and red arrowheads, respectively. The predicted pseudoduplex length and the maximum MutSβ bound are shown below EMSA gels. **c,** The cryoEM maps of the ATP-binding pockets in MSH2 and MSH3 in the 1-us (CAG)n stem structure. MSH2 with ΔN MSH3, but not FL MSH3, contains an ADP, which remained bound throughout the protein purification as observed previously[2].

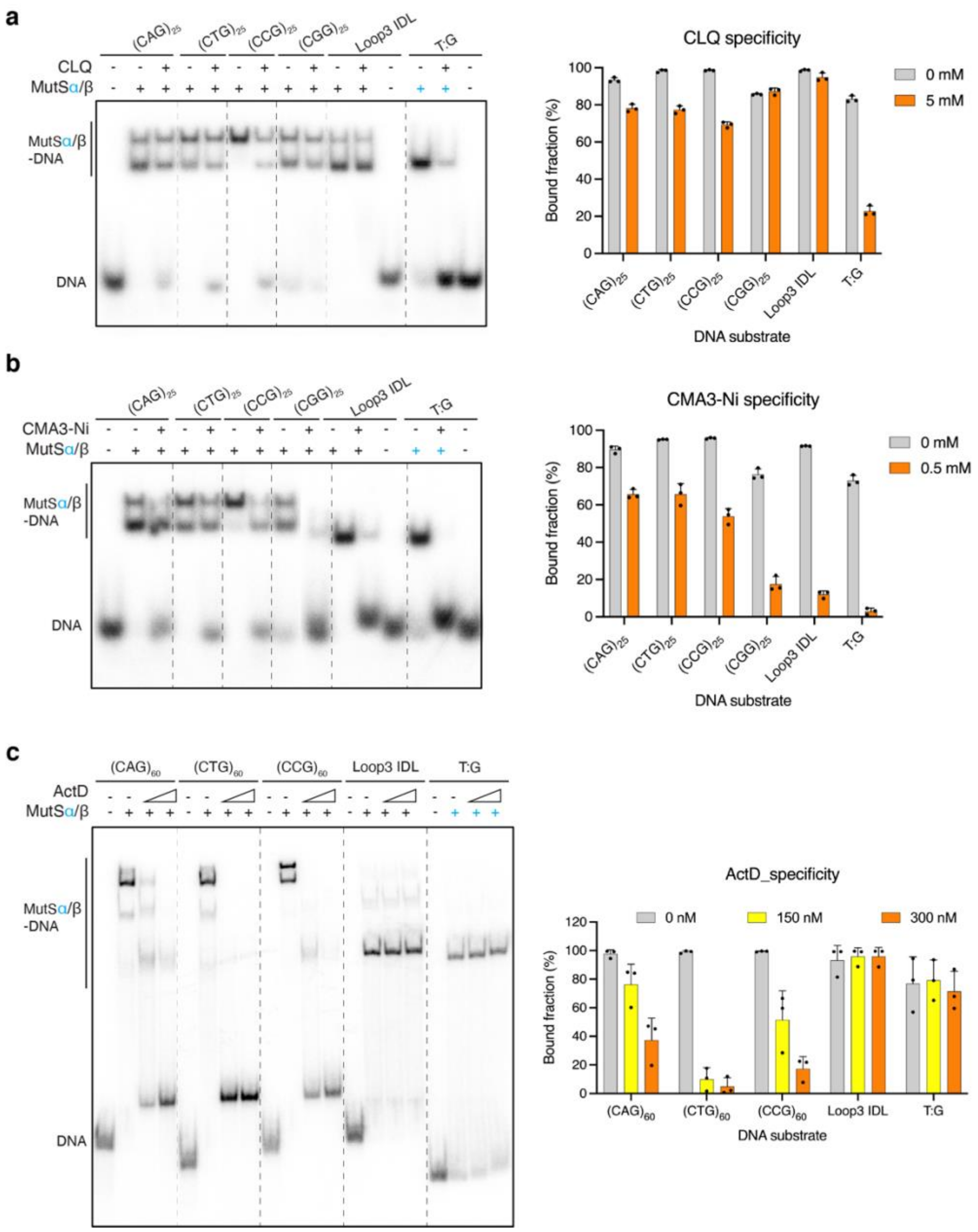


**Extended Data Fig. 8. Inhibition of MutSβ binding to CNG repeats by DNA intercalators**

**a-b,** The inhibition effects of CLQ (**a**) and CMA3-Ni (**b**) on MutSβ binding to $(CNG)_{25}$ (N=A, T, C, G) or MutS (α or β) binding T:G or IDL mismatches of similar length. **c,** The inhibition effects of ActD on MutSβ binding to $(CNG)_{60}$ (N=A, T, C) or MutS (α or β) binding T:G or IDL mismatches of similar length. Average of triplicate measurements is shown with the standard deviations in the bar graphs.

### Extended Data Table 1. Cryo-EM data collection, refinement and validation statistics

| | MutS$\beta$ complexed with $(CAG)_{25}$ and d40h30 | | | MutS$\beta$ complexed with $(CTG)_{25}$ | | | MutS$\beta$ complexed with uneven bubble | |
|---|---|---|---|---|---|---|---|---|
| | 1-us stem | 2-us stem | Hairpin end | 1-us stem | 2-us stem | Hairpin end | $(CAG)_2$/CAG | $(CAG)_2/T_3$ |
| **EMDB** | 42359 | 42360 | 42361 | 42362 | 42367 | 42368 | 42369 | 42370 |
| **PDB** | 8ULL | 8ULO | 8ULP | 8ULQ | 8ULV | 8ULW | 8ULX | 8ULY |
| **Data collection and processing** | | | | | | | | |
| Magnification | | 105,000 | | | 105,000 | | 45,000 | 45,000 |
| Voltage (kV) | | 300 | | | 300 | | 200 | 200 |
| Electron exposure (e–/Å$^2$) | | 49.9 | | | 48.2 | | 70.0 | 70.0 |
| Defocus range (μm) | | -0.5 to -1.5 | | | -0.5 to -1.5 | | -0.5 to -1.5 | -0.5 to -1.5 |
| Pixel size (Å) | | 0.4165 | | | 0.4165 | | 0.4340 | 0.4340 |
| Symmetry imposed | C1 | C1 | C1 | C1 | C1 | C1 | C1 | C1 |
| Initial particle images (no.) | 1,741,059 | 1,741,059 | 1,741,059 | 1,499,795 | 1,499,795 | 1,499,795 | 875,183 | 3,734,363 |
| Final particle images (no.) | 136,035 | 9,214 | 53,925 | 51,944 | 136,195 | 17,296 | 227,747 | 239,714 |
| Map resolution (Å, FSC=0.143) | 2.8 | 3.4 | 3.0 | 3.1 | 2.9 | 3.4 | 3.2 | 3.2 |
| **Refinement** | | | | | | | | |
| Initial model used (PDB code) | 3THX | 8ULL | 8ULL | 8ULL | 8ULL | 8ULL | 8ULL | 8ULL |
| CC_mask | 0.89 | 0.88 | 0.90 | 0.89 | 0.89 | 0.89 | 0.88 | 0.88 |
| CC_volume | 0.88 | 0.87 | 0.89 | 0.88 | 0.88 | 0.88 | 0.87 | 0.87 |
| CC_peaks | 0.78 | 0.78 | 0.79 | 0.79 | 0.79 | 0.80 | 0.79 | 0.77 |
| Model resolution (Å, FSC=0.5) | 2.9 | 3.4 | 3.0 | 3.2 | 3.0 | 3.4 | 3.3 | 3.3 |
| Map sharpening *B* factor (Å$^2$) | -81.3 | -58.4 | -70.4 | -85.4 | -92.5 | -71.0 | -91.5 | -89.3 |
| Model composition | | | | | | | | |
| Non-hydrogen atoms | 14,954 | 14,969 | 14,538 | 15,023 | 15,052 | 14,601 | 15,049 | 15,096 |
| Protein residues | 1,760 | 1,753 | 1,761 | 1,760 | 1,755 | 1,762 | 1,761 | 1,759 |
| Nucleotides | 45 | 48 | 24 | 45 | 48 | 27 | 49 | 49 |
| Ligands | 0 | 0 | 0 | 0 | 0 | 0 | 0 | 0 |
| *B* factors (Å$^2$) | | | | | | | | |
| Protein | 134.33 | 112.64 | 126.15 | 134.25 | 140.16 | 125.72 | 108.47 | 132.80 |
| Nucleotides | 253.39 | 236.18 | 189.61 | 195.81 | 201.02 | 179.53 | 152.08 | 185.80 |
| Ligand | 0 | 0 | 0 | 0 | 0 | 0 | 0 | 0 |
| R.m.s. deviations | | | | | | | | |
| Bond lengths (Å) | 0.002 | 0.003 | 0.003 | 0.003 | 0.003 | 0.003 | 0.003 | 0.003 |
| Bond angles (°) | 0.510 | 0.557 | 0.577 | 0.585 | 0.595 | 0.566 | 0.564 | 0.572 |
| Validation | | | | | | | | |
| MolProbity score | 1.46 | 1.59 | 1.60 | 1.61 | 1.68 | 1.54 | 1.53 | 1.47 |
| Clashscore | 7.12 | 10.69 | 8.06 | 9.63 | 7.85 | 8.96 | 9.39 | 8.83 |
| Poor rotamers (%) | 1.22 | 0.26 | 1.35 | 0.32 | 1.61 | 0.26 | 0.13 | 0.26 |
| Ramachandran plot | | | | | | | | |
| Favored (%) | 98.1 | 97.8 | 97.7 | 97.5 | 97.5 | 97.7 | 97.9 | 98.3 |
| Allowed (%) | 2.0 | 2.2 | 2.3 | 2.5 | 2.5 | 2.3 | 2.1 | 1.7 |
| Disallowed (%) | 0.0 | 0.0 | 0.0 | 0.0 | 0.0 | 0.0 | 0.0 | 0.0 |

### Extended Data Table 2. DNA oligos used in cryo-EM and binding studies

| DNA oligo | Sequence | Substrate | Fig. # * |
|---|---|---|---|
| Loop4_IDL_F | TCATCGATCGCACAGCTTCAGATAGG | Loop4 IDL | 1b, ED 1a |
| Loop4_IDL_R | CCTCTATCTGAAGCCGATCGATGG | | |
| CAG8 | CCCAGCAGCAGCAGCAGCAGCAGCAGGG | $(CAG)_8$ | 1b, ED 1a, 7b |
| CAG18 | CCCAGCAGCAGCAGCAGCAGCAGCAGCAGCAGCAGCAGCAGCAGCAGCAGCAGCAG<br>GG | $(CAG)_{18}$ | 1b, ED 1a, 4c,d |
| CAG25 | CAGCAGCAGCAGCAGCAGCAGCAGCAGCAGCAGCAGCAGCAGCAGCAGCAGCAGCA<br>GCAGCAGCAGCAGCAGCAG | $(CAG)_{25}$ | 1c,d, ED 1b, 2, 7b |
| CTG25 | CTGCTGCTGCTGCTGCTGCTGCTGCTGCTGCTGCTGCTGCTGCTGCTGCTGCTGCT<br>GCTGCTGCTGCTGCTG | $(CTG)_{25}$ | 1c,d, ED 1b, 2 |
| CCG25 | CCGCCGCCGCCGCCGCCGCCGCCGCCGCCGCCGCCGCCGCCGCCGCCGCCGCCGC<br>CGCCGCCGCCGCCGCCG | $(CCG)_{25}$ | 1c,d, ED 1b |
| CGG25 | CGGCGGCGGCGGCGGCGGCGGCGGCGGCGGCGGCGGCGGCGGCGGCGGCGGCG<br>GCGGCGGCGGCGGCGGCGG | $(CGG)_{25}$ | 1c,d, ED 1b |
| GAA25 | GAAGAAGAAGAAGAAGAAGAAGAAGAAGAAGAAGAAGAAGAAGAAGAAGAAGAAG<br>AAGAAGAAGAAGAAGAAGAA | $(GAA)_{25}$ | 1c,d, ED 1b, 6 |
| LoopCAG2 | GACGCTGACACTGGTGCTTGCAGCAGGCAGCTGTCGTGGCGAGTGC | $(CAG)_2$/CAG and $(CAG)_2/T_3$ uneven bubble | ED 4 |
| compLoopCAG | GCACTCGCCACGACAGCTGCCAGCAAGCACCAGTGTCAGCGTC | $(CAG)_2$/CAG uneven bubble | ED 4 |
| compLoopT3 | GCACTCGCCACGACAGCTGCTTTCAAGCACCAGTGTCAGCGTC | $(CAG)_2/T_3$ uneven bubble | ED 4 |
| CTG60 | CTGCTGCTGCTGCTGCTGCTGCTGCTGCTGCTGCTGCTGCTGCTGCTGCTGC<br>TGCTGCTGCTGCTGCTGCTGCTGCTGCTGCTGCTGCTGCTGCTGCTGCTGCT<br>GCTGCTGCTGCTGCTGCTGCTGCTGCTGCTGCTGCTGCTGCTGCTGCTGCTG<br>CTGCTGCTGCTGCTGCTGCTGCTG | $(CTG)_{60}$ | 4a,b, ED 7a, 8c |
| CAG60 | CAGCAGCAGCAGCAGCAGCAGCAGCAGCAGCAGCAGCAGCAGCAGCAGCAGCAGCA<br>GCAGCAGCAGCAGCAGCAGCAGCAGCAGCAGCAGCAGCAGCAGCAGCAGCAGCAGC<br>AGCAGCAGCAGCAGCAGCAGCAGCAGCAGCAGCAGCAGCAGCAGCAGCAGCAGCAG<br>CAGCAGCAGCAG | $(CAG)_{60}$ | 4a-d, ED 7b, 8c |
| CCG60 | CCGCCGCCGCCGCCGCCGCCGCCGCCGCCGCCGCCGCCGCCGCCGCCGCCGCCGC<br>CGCCGCCGCCGCCGCCGCCGCCGCCGCCGCCGCCGCCGCCGCCGCCGCCGCCGCC<br>GCCGCCGCCGCCGCCGCCGCCGCCGCCGCCGCCGCCGCCGCCGCCGCCGCCGCC<br>GCCGCCG | $(CCG)_{60}$ | ED 8c |
| CAG30 | CAGCAGCAGCAGCAGCAGCAGCAGCAGCAGCAGCAGCAGCAGCAGCAGCAGCAGCA<br>GCAGCAGCAGCAGCAGCAGCAGCAGCAGCAG | $(CAG)_{30}$ | 4c,d |
| CAG45 | CAGCAGCAGCAGCAGCAGCAGCAGCAGCAGCAGCAGCAGCAGCAGCAGCAGCAGCA<br>GCAGCAGCAGCAGCAGCAGCAGCAGCAGCAGCAGCAGCAGCAGCAGCAGCAGCAGC<br>AGCAGCAGCAGCAGCAGCAGCAG | $(CAG)_{30}$ | 4c,d |
| Loop3_IDL_F | GACGCTGACACTGGTGCTTGCAGCAGGCAGCTGTCGTGGCGAGTGC | Loop3 IDL | 4c,d, ED7a |
| Loop3_IDL_R | GCACTCGCCACGACAGCTGCCTGCAAGCACCAGTGTCAGCGTC | | |
| TriArm_0 | GCAACCTATAGGACGCTGACACTGGTGCTGGCAGCTGTCGTGGCGAGTGCAGTGCT | TriArm_9-9, TriArm_9-6, TriArm | ED 1c-e |
| TriArm_1 | AGCACTGCACTCGCCACGACAGCTGCCACAGCAGCAGGTCAGCTTCAAGTGGATCGG | TriArm_9-9, TriArm_9-6, TriArm | ED 1c-e |
| TriArm_2 | CCGATCCACTTGAAGCTGACCAGCAGCAGGCACCAGTGTCAGCGTCCTATAGGTTGC | TriArm_9-9 | ED 1c,d |
| TriArm_3 | CCGATCCACTTGAAGCTGACCAGCAGGCACCAGTGTCAGCGTCCTATAGGTTGC | TriArm_9-6 | ED 1c-e |
| TriArm_4 | CCGATCCACTTGAAGCTGACCTGCTGCTGGCACCAGTGTCAGCGTCCTATAGGTTGC | TriArm | ED 1c,d |
| CAG30_40-40 | CCATGTCAAGCGAGCACTGCACTCGCCACGACAGCTGCCACAGCAGCAGCAGCAGCA<br>GCAGCAGCAGCAGCAGCAGCAGCAGCAGCAGCAGCAGCAGCAGCAGCAGCAGCAGC<br>AGCAGCAGCAGCAGCAGGCACCAGTGTCAGCGTCCTATAGGTTGCACCATCGTTCCA | d40h30 | ED 2 |
| compCAG_40-40 | TGGAACGATGGTGCAACCTATAGGACGCTGACACTGGTGCTGGCAGCTGTCGTGGCG<br>AGTGCAGTGCTCGCTTGACATGG | | |
| T21-21 | ATAGGACGCTGACACTGGTGCTTGGCAGCTGTCGTGGCGAGTG | TG_MM_42bp | 4e,f, ED 8a,b |
| G21-21 | CACTCGCCACGACAGCTGCCAGGCACCAGTGTCAGCGTCCTAT | | |
| T40-40 | TGGAACGATGGTGCAACCTATAGGACGCTGACACTGGTgCtTGGCAGCTGTCGTGGCG<br>AGTGCAGTGCTCGCTTGACATGG | TG_MM_80bp | ED 8c |
| G40-40 | CCATGTCAAGCGAGCACTGCACTCGCCACGACAGCTGCCAgGcACCAGTGTCAGCGTC<br>CTATAGGTTGCACCATCGTTCCA | | |
| Loop3_CAG_20-20 | CATCGCCACGACAGCTGCCACAGGcACCAGTGTCAGCGTTACC | Loop3_IDL_40bp | 4e,f, ED 8a,b |
| Loop3_comp_20-20 | GGTAACGCTGACACTGGTgCTGGCAGCTGTCGTGGCGATG | | |
| Loop3_CAG_45-45 | GGCATTGTCATTGCTCATGTAGCCTCAAACTCCTGGGCTCAAGCCCAGATCCTCCTGC<br>CTTAGCTTCCTAGTAGCAAGGACCACAGGTGCATG | Loop3_IDL_90bp | ED 8c |
| Loop3_comp_45-45 | CATGCACCTGTGGTCCTTGCTACTAGGAAGCTAAGGCAGGAGGATGGCTTGAGCCCA<br>GGAGTTTGAGGCTACATGAGCAATGACAATGCC | | |

* ED: Extended Data Fig.
